# Cophenetic Spatial Topology Embedding reveals multiscale tissue architecture in spatial omics

**DOI:** 10.64898/2026.05.26.727847

**Authors:** Mengping Long, Taobo Hu, Alexandros Sountoulidis, Christos Samakovlis, Mats Nilsson

**Author notes:** Correspondance: Mengping Long,; Taobo Hu,; Mats Nilsson. These authors contributed equally to this work (Mengping Long, Taobo Hu).

## Abstract

The spatial organization of tissues emerges from cell interactions across multiple scales, yet current spatial omics analysis tools often emphasize local neighborhoods and may not summarize broader tissue architecture. Here we introduce Cophenetic Spatial Topology Embedding (COSTE), a computational framework that embeds directed nearest-neighbor distance profiles into a hierarchical metric space without requiring the user to define a spatial radius or neighborhood cutoff. COSTE can be applied to cell-level and single-transcript inputs without requiring cell segmentation. It constructs directed distance profiles between cell populations and uses hierarchical clustering to quantify tissue topology. This yields a Spatial Separation Score (SSS), a sample-normalized score from 0 to 1 that summarizes relative spatial separation within an analyzed tissue. We apply COSTE to spatial transcriptomics datasets of pulmonary fibrosis and triple-negative breast cancer (TNBC), where it delineates tissue structures, nominates spatially defined cell states, and highlights disease-or treatment-associated architectural patterns that are not readily captured by local neighborhood-based analyses. Our approach provides an interpretable framework for exploring tissue architecture and cell-cell spatial relationships in spatial omics data.

## Introduction

Recent *in situ* spatial transcriptomics technologies (such as 10x *Xenium*^1, 2^, RIBOmap^3^, NanoString *CosMx*^4^, and Vizgen *MERFISH*^5, 6^) now enable gene expression mapping within intact tissues at subcellular resolution, offering unprecedented opportunities to study how diverse cell types organize into functional structures. However, current computational tools struggle to fully exploit these rich datasets. Many methods have limited scalability or require sensitive parameter tuning^7^, and they often infer spatial groupings that do not correspond to known histological units (e.g., failing to recover an anatomic structure like a breast lobule or a lung airway)^8^, which in turn undermines biological interpretability. There is a clear need for a **scalable**, **generalizable**, and **segmentation-free** analytical framework to chart tissue organization from spatial omics data^9^.

Most existing approaches – whether classical or machine-learning–based – rely on analyzing local cell neighborhoods (for example, niche enrichment or nearest-neighbor graph methods)^10–13^. These techniques typically assess each cell’s composition within a fixed radius or *k*-nearest neighbors^14^. While effective for detecting short-range cell–cell interactions, such local metrics inherently overlook long-range relationships and constrain the analysis to a single predefined spatial scale^15^. Yet histopathology has long shown that tissues exhibit **hierarchical organization across multiple scales** – from microscale cellular interactions to macroscale tissue architecture^16^.

For instance, at a larger spatial scale, a tumor can be broadly delineated from adjacent normal tissue, whereas at a finer scale the tumor region contains intermixed tumor cells and stromal cells. Crucially, interactions **across** these scales – such as whether tumor cells infiltrate into surrounding normal tissue or instead push against it to form an expansively growing mass – reflect important biological properties of the tumor^17^. These cross-scale patterns are often missed by current neighborhood-based frameworks. Moreover, the “micro-niches” identified by local analyses are difficult to clearly define, quantify, or interpret in anatomical terms, complicating downstream biological insight^18, 19^.

To overcome these challenges, we developed Cophenetic Spatial Topology Embedding (COSTE), a computational framework that embeds multiscale spatial relationships into a hierarchical metric space without requiring a user-defined spatial radius or neighborhood cutoff. COSTE can be applied to both cell-type and single-transcript inputs without requiring cell segmentation. It constructs directed nearest-neighbor distance profiles between all pairs of cell types and then performs hierarchical clustering to summarize tissue spatial topology in an ultrametric tree. From this clustering, we derive a Spatial Separation Score (SSS) between each pair of cell populations, normalized from 0 to 1 within each analyzed sample, which measures their relative separation in the COSTE hierarchy.

Because COSTE evaluates the arrangement of all annotated cell types together, it summarizes both short-range and longer-range spatial relationships in one analysis. It does not require the user to predefine a distance cutoff or neighborhood size, and by averaging directed cross-type distances at the cell-type level it reduces the dominance of local neighbor-count imbalance. At the same time, nearest-neighbor distances remain influenced by local density and sampling, so SSS values should be interpreted as relative spatial patterns generated under a shared analysis workflow rather than as absolute physical distances. By comparing spatial distance profiles across all cell types, COSTE can nominate biologically interpretable structures-within-structures - groups of cell types that co-localize to form candidate functional niches.

We demonstrate the capabilities of COSTE across diverse spatial omics datasets, including a whole neonatal mouse tissue, a pulmonary fibrosis spatial transcriptomics cohort^23^, and a spatial proteomics dataset from a human triple-negative breast cancer (TNBC) trial^24^. In these applications, COSTE delineates global tissue architecture, nominates spatially confined cell subpopulations and niches, and highlights disease- or treatment-associated architectural patterns that complement local neighborhood-based analyses. By providing a quantitative, hierarchical view of tissue organization, COSTE offers a general and interpretable approach to explore complex spatial patterns and cell-cell relationships in tissues.

## Results

### 1. A global, hierarchical representation of tissue spatial topology

COSTE embeds the spatial architecture of a tissue by comparing directed nearest-neighbor distance profiles between all cell-type pairs (Fig. 1a). Each cell type’s spatial pattern is represented by its distances to every other cell type. These directed inter-type distances are aggregated into a matrix and subjected to hierarchical clustering, producing a dendrogram in which cell types with similar spatial profiles cluster together. We then compute the Spatial Separation Score (SSS) for each pair of cell types from the cophenetic distances in the searcher-based dendrogram, normalizing the scores between 0 and 1 within each analyzed sample. The resulting SSS heatmap provides a global view of tissue organization: cell types that co-localize in space form nested low-SSS blocks in the heatmap, whereas more separated cell-type pairs appear as high-SSS regions (Fig. 1b). The hierarchy encoded by the dendrogram allows us to inspect multiple levels of spatial structure by cutting the dendrogram at different heights, yielding coarse to fine-grained groupings of cell types.

**Figure 1.**
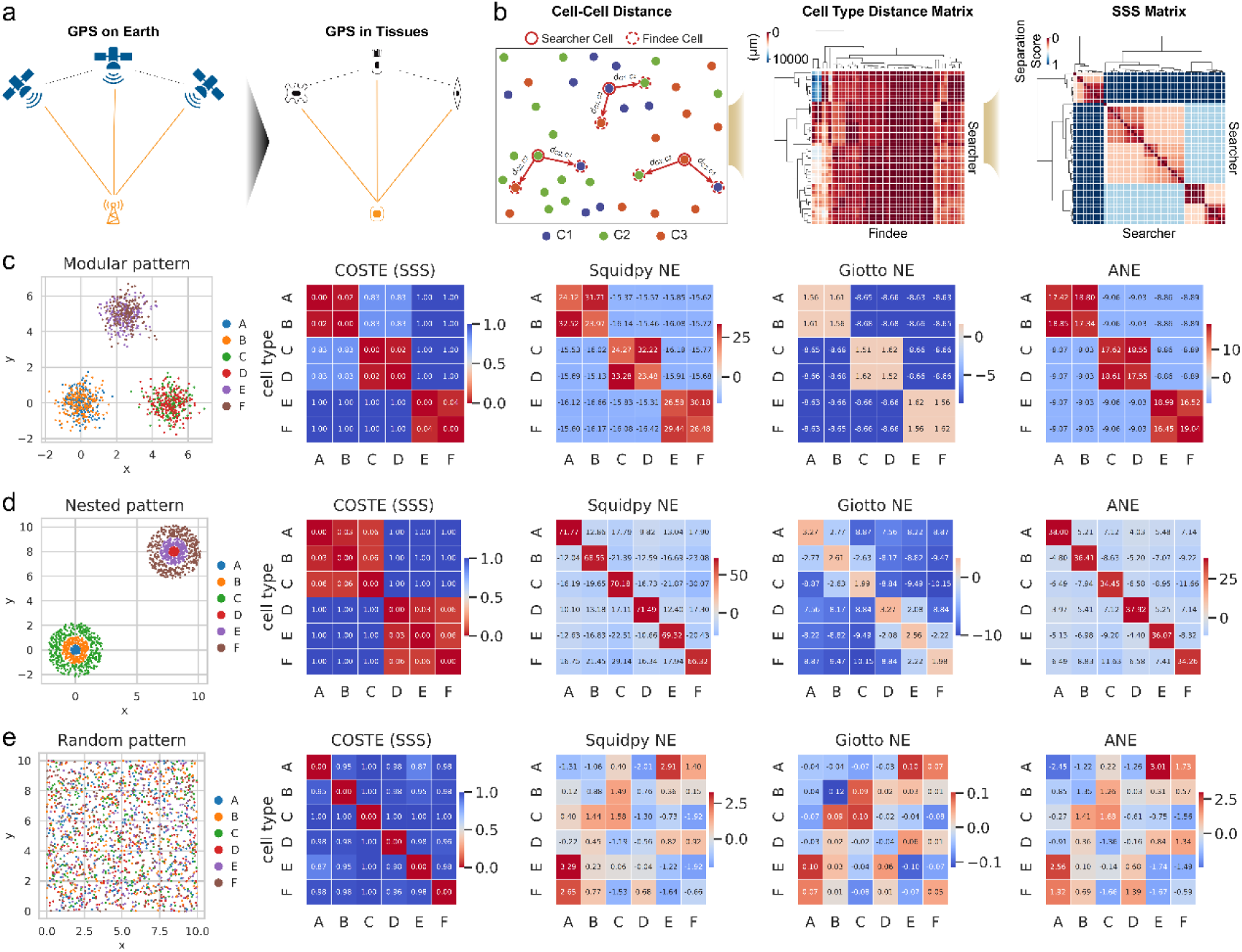
Conceptual overview of the COSTE framework and benchmarking on synthetic datasets. (a) Conceptual motivation for representing each cell type by its directed nearest-neighbor distance profile to other cell types. (b) Schematic of the COSTE pipeline. For every searcher cell type, nearest-neighbor distances to all findee cell types are computed and averaged into a directed inter-type distance matrix. Row and column profiles can be clustered separately, and the searcher-based dendrogram is used to compute normalized cophenetic distances, yielding the Spatial Separation Score (SSS). (c-e) Synthetic benchmarks illustrating how COSTE and local neighborhood-enrichment baselines summarize known spatial architectures. (c) Modular pattern consisting of three well-separated co-localized cell-type pairs. (d) Nested pattern in which one cell-type cluster is embedded within another. (e) Random pattern with no true spatial structure. COSTE captures the modular and nested patterns in the SSS representation, while all methods correctly report the absence of structure in the random pattern.

We first evaluated COSTE on synthetic spatial datasets and compared it with three local neighborhood-enrichment baselines: Squidpy NE^13^, Giotto NE^10^, and Analytical Neighborhood Enrichment (ANE)^25^. The benchmarks comprised a modular pattern with three distinct spatial blocks (Fig. 1c), a nested pattern in which one cell-type cluster is enclosed within another (Fig. 1d), and a random pattern with no true spatial structure (Fig. 1e). These tests assessed how each representation summarized known spatial architecture. In the modular dataset, all four approaches produced block-like association patterns consistent with the expected domains. In the nested configuration, COSTE’s hierarchical representation separated the two nested modules as structure-within-structure patterns in the SSS heatmap, whereas the local neighborhood-enrichment baselines did not recover the full nested arrangement under the tested settings. We next generated twelve nested patterns with systematic perturbations in shell spacing, shell thickness and inter-group distance (Supplementary Fig. 2). Across these variants, COSTE preserved the two nested structures in the SSS representation (Supplementary Fig. 3), while Squidpy (Supplementary Fig. 4), Giotto (Supplementary Fig. 5) and ANE (Supplementary Fig. 6) showed more limited separation of these hierarchical geometries under the tested parameter settings. Lastly, all methods appropriately found no spurious co-localization in the random dataset. These simulations illustrate a setting in which a global hierarchical representation captures nested geometry more directly than local neighborhood-enrichment summaries; they are not intended as an exhaustive head-to-head benchmark against all methods designed for hierarchical or multiscale spatial niche discovery.

We next applied COSTE to a spatial transcriptomics atlas of a neonatal mouse pup (44 cell types profiled by Xenium). Without specifying a neighborhood radius, COSTE organized the tissue into a structure-within-structure hierarchy of spatial relationships (Fig. 2a and Supplementary Fig. 7). This unsupervised analysis recapitulated several known anatomical structures in the mouse. For example, all cell types belonging to the retina clustered into one top-level branch of the dendrogram, consistent with the retina’s stratified anatomy. Another branch encompassed enteric neurons, which were segregated from the various central nervous system cell types. COSTE also highlighted more subtle spatial partitions: for instance, two transcriptionally distinct fibroblast subtypes - Thbs2+Tnxb+ fibroblasts and Thbs4+Mfap5+ fibroblasts - were placed into different spatial structures (S3 and S4; Fig. 2d), consistent with their distinct locations (S3 fibroblasts were found mainly in the head and tail regions, whereas S4 fibroblasts were enriched in the trunk/abdominal region). The local neighborhood-enrichment baselines suggested weaker separation of these fibroblast subtypes (Supplementary Fig. 8), whereas the COSTE embedding provided a global view of this tissue substructure.

**Figure 2.**
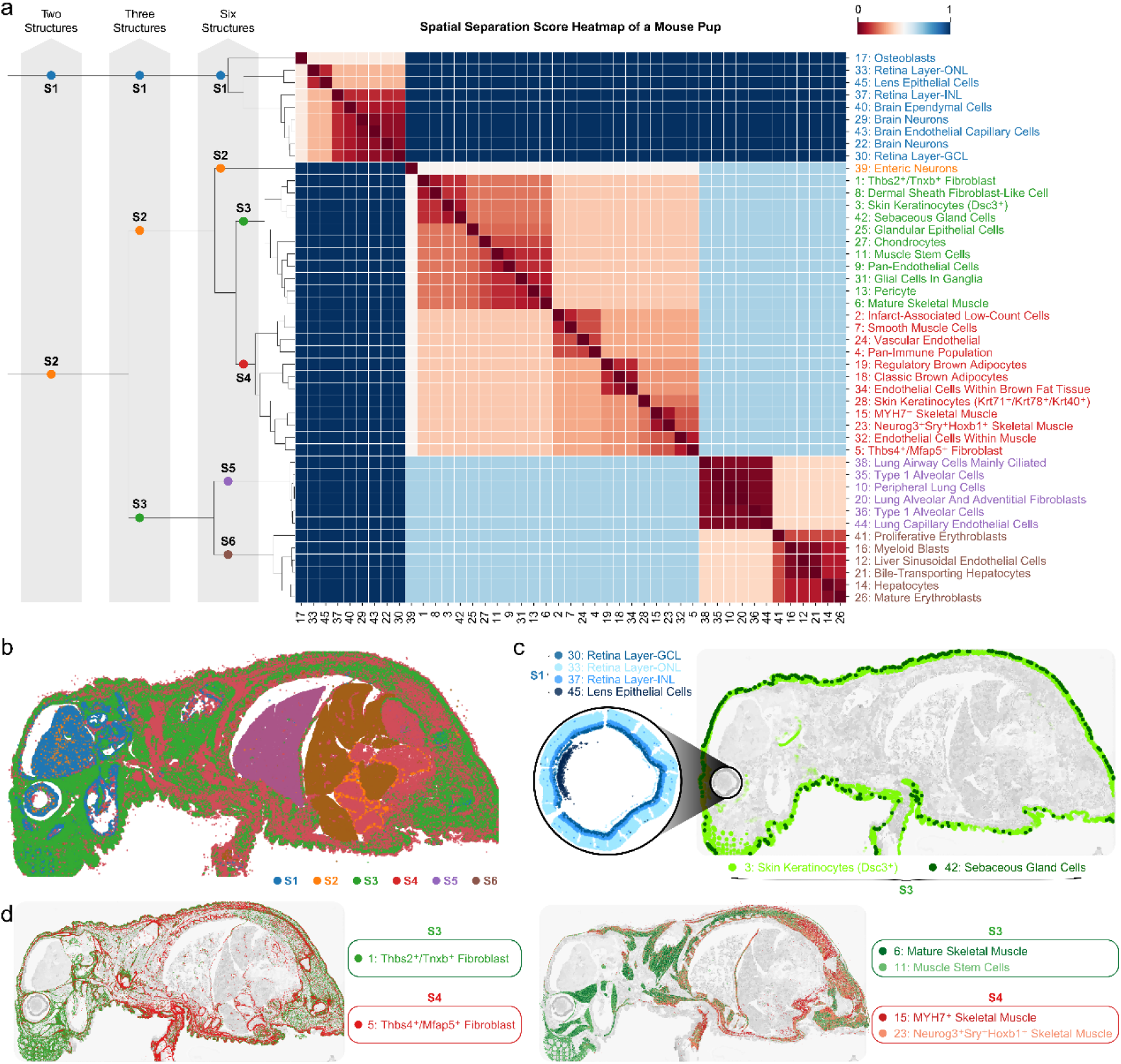
Overview of multiscale spatial mapping by COSTE in a neonatal mouse tissue. (a) SSS heatmap for an in situ transcriptomics dataset of a neonatal mouse (44 cell types, 10x Xenium platform) summarizes spatial organization across multiple scales. Cutting the dendrogram at different levels yields 2, 3, or 6 nested spatial structures (S1-S6), with red blocks indicating cell-type groups that co-localize and blue blocks indicating groups that are spatially segregated. (b) Spatial mapping of selected cell types illustrates these multiscale structures. (c) The three retinal layers (ganglion cell layer, inner nuclear layer, outer nuclear layer) form a tightly interacting unit (structure S1), and skin keratinocytes co-localize with sebaceous gland cells (also part of S1). (d) Two fibroblast subtypes (Thbs2+Tnxb+ versus Thbs4+Mfap5+) and two groups of skeletal muscle cells separate into distinct domains (structures S3 vs. S4), illustrating spatial patterns captured by the global hierarchical representation and not emphasized by local-neighborhood summaries.

We further assessed whether local neighborhood-enrichment summaries could recover longer-range spatial relationships by varying their radius and number-of-neighbors parameters. In the tested settings, no single neighborhood definition reproduced all of the global structures observed in the COSTE hierarchy. For example, the mouse retina’s layered organization was most apparent in the COSTE representation (Fig. 2c), whereas Squidpy NE produced parameter-sensitive groupings across six tested settings (Supplementary Fig. 9). COSTE also highlighted spatial proximity between keratinocytes and rare sebaceous gland cells in skin tissue (Fig. 2d), a relationship that local neighbor counts can underweight when abundant keratinocytes primarily neighbor themselves (Supplementary Fig. 9). This example illustrates that directed cross-type distance profiles can reduce the dominance of abundant self-neighbor counts, although the underlying nearest-neighbor distances remain sensitive to local density and sampling. Together, these results show that COSTE provides a complementary hierarchical representation of tissue topology, capturing anatomical arrangements and candidate spatial partitions in a complex tissue.

Finally, we benchmarked the computational efficiency of COSTE. We recorded detailed runtime and memory usage statistics for the above mouse pup analysis, including total CPU time, peak memory consumption (resident set size, RSS), minor page faults, context switches, and processing throughput (cells analyzed per second). The results, compiled in Supplementary Table 1, revealed major differences in scalability. COSTE and ANE completed the ∼1 million-cell analysis with lower run time and memory usage in this benchmark, reflecting their avoidance of costly random permutations. In contrast, Squidpy and Giotto - both of which rely on permutation sampling for neighborhood enrichment - required more computation and memory under the tested configuration. This highlights the computational efficiency of COSTE for large cell-by-coordinate datasets.

### 2. COSTE maps multiscale spatial organization and spatial niches

To benchmark COSTE on a tissue with known discrete compartments, we applied it to a human lymph node section (28 annotated cell types profiled by Xenium). The COSTE analysis highlighted three spatial structures corresponding to canonical functional niches of lymph node anatomy^26^: an outer cortex dominated by B cells, an inner medulla enriched in plasma cells, macrophages, stromal and endothelial cells, and a paracortex region populated mainly by T cells (Supplementary Fig. 10). The inferred groupings were consistent with prior histological knowledge of this tissue. This result illustrates how a global hierarchical representation can summarize tissue substructures that mirror known biology, including both broad lymph-node regions and more specific cell-composition-defined niches within those regions.

### 3. Identifying spatial remodeling in disease progression

We next examined whether COSTE could help interpret changes in tissue architecture during disease progression. To this end, we analyzed a Xenium spatial transcriptomics pulmonary fibrosis cohort consisting of 45 tissue samples grouped into healthy, mildly affected (less affected), and severely affected (more affected) stages^23^. Using a shared cell-type annotation and analysis workflow, the average SSS heatmap of healthy lung samples delineated modules corresponding to known lung structures, including an airway structure and a terminal respiratory unit (TRU) structure (the gas-exchanging alveolar region) (Fig. 3a and 4a). In fibrotic lungs, these modules were less ordered. COSTE showed increased sample-normalized SSS between alveolar epithelial cells (AT1/AT2) and capillary endothelial cells, consistent with weaker relative coupling of alveoli and capillaries during fibrosis. We summarized this pattern using a TRU Remodeling Score (TRS), defined as the within-sample SSS between AT2 cells and capillary cells. TRS should be interpreted as a relative architectural index computed under the same workflow, not as an absolute distance that is intrinsically comparable across all tissues. Within this cohort, TRS showed an ordered shift from healthy to more severely fibrotic lungs (Fig. 3b,c). A localized analysis of TRS in histologically annotated regions was consistent with this interpretation: regions of pronounced fibrosis showed higher TRS than less affected areas (Fig. 3d).

**Figure 3.**
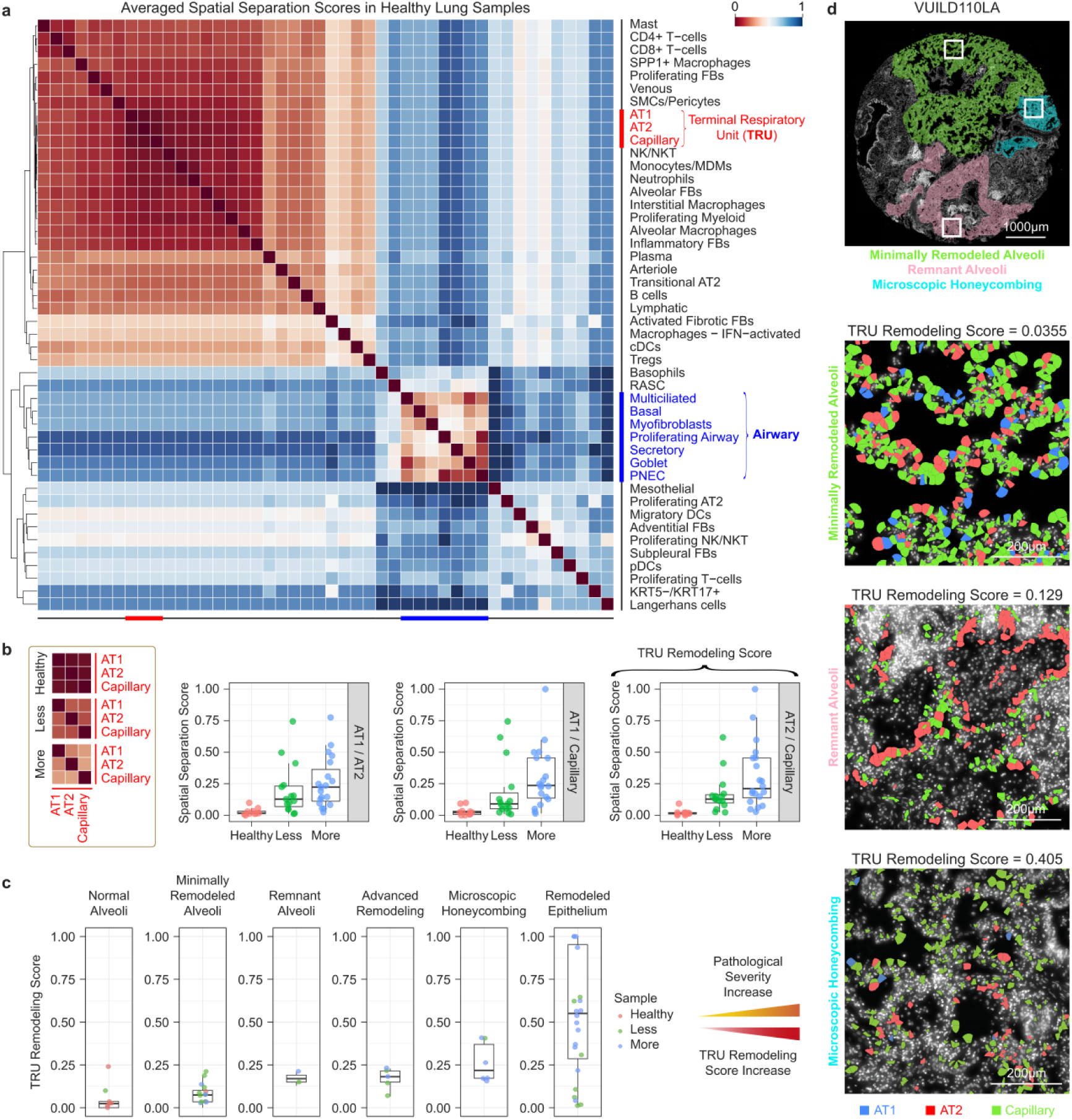
COSTE summarizes spatial remodeling of lung alveolar structures in pulmonary fibrosis. (a) In healthy lungs, COSTE’s Spatial Separation Score (SSS) heatmap delineates two compartments - the gas-exchanging Terminal Respiratory Unit (TRU) and the airway - each showing internal co-localization and separation from one another. (b) Across fibrosis severity groups analyzed with the same workflow, the sample-normalized SSS between alveolar epithelial cells (AT1, AT2) and capillary endothelial cells increases, consistent with weaker relative alveolar-capillary coupling. (c) The SSS between AT2 and capillary cells, defined as the TRU Remodeling Score (TRS), shows an ordered shift from healthy to severely fibrotic lungs and should be interpreted as a relative architectural index. (d) In a representative fibrotic sample, regions classified as minimally remodeled alveoli, remnant alveoli, and microscopic honeycombing display increasing TRS values, visualized by local uncoupling of AT1 (blue), AT2 (red), and capillary (green) cells in those areas. These patterns illustrate how COSTE summarizes progressive TRU architectural disruption during pulmonary fibrosis.

**Figure 4.**
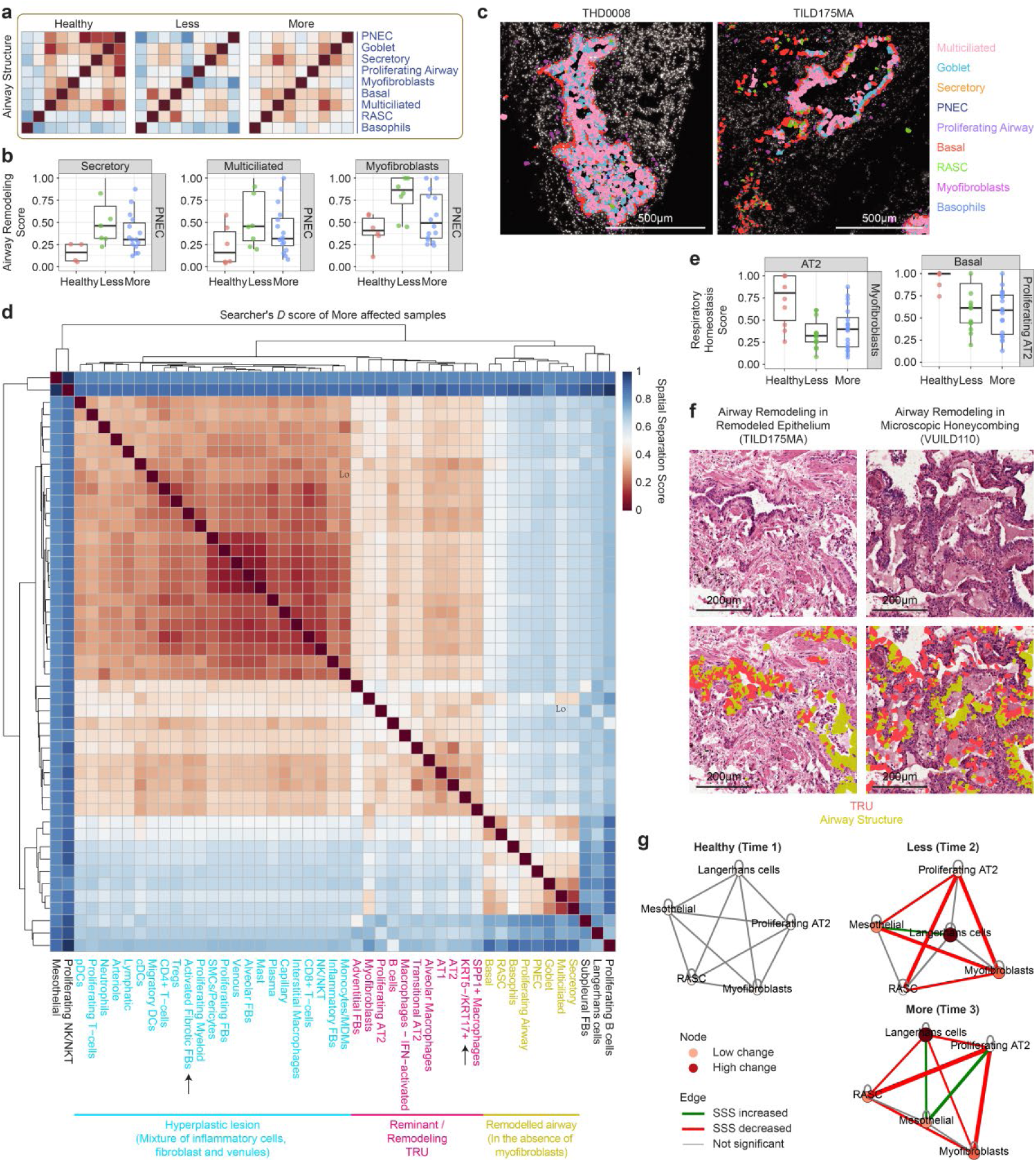
COSTE and exploratory DST-GNN summarize pathological structural reorganization in fibrotic lungs. (a-c) In pulmonary fibrosis, COSTE’s SSS analysis shows that airway-associated cell types become increasingly separated from one another across disease stages, consistent with progressive disruption of normal airway architecture. (d) In severely affected lungs, the method highlights candidate spatial structures: hyperplastic lesions enriched in inflammatory cells, fibroblasts and venules; remnant TRU-like clusters containing proliferating AT2 and basaloid cells; and remodeled airway regions devoid of myofibroblasts. (e, f) Correspondingly, the spatial separation between selected cell pairs drops in late-stage disease - for example, AT2 epithelial cells and myofibroblasts, as well as basal cells and proliferating AT2 cells, become more intermingled - consistent with abnormal mixing of alveolar (TRU) and airway compartments. Histological images support this structural interpretation in fibrotic lesions. (g) An exploratory diffusion-based spatio-temporal graph neural network (DST-GNN) applied to sequential COSTE graphs prioritizes candidate interactions involving proliferating AT2 cells, respiratory airway secretory cells (RASC), and myofibroblasts; these are interpreted as hypotheses for follow-up rather than validated drivers.

To visualize how normal lung architecture changes in fibrosis, we generated unclustered SSS heatmaps by fixing the row/column order to that of the healthy lung hierarchy and projecting fibrotic samples onto it. This visualization suggested loss of the ordered airway-TRU structure in diseased lungs and the emergence of alternative groupings of cell types (Supplementary Fig. 11). COSTE further highlighted candidate pathological structures in severely affected lungs (Fig. 4b-4f). For example, in advanced fibrosis the analysis separated hyperplastic lesions (clusters of proliferative epithelial and myofibroblast-rich areas) from residual, partially intact alveolar units. These findings illustrate how a global spatial embedding can summarize complex remodeling events that involve the reorganization of multiple cell types.

Beyond static descriptions, we explored whether temporal graph modeling could prioritize candidate cell-cell relationships associated with these architectural changes. We therefore applied a Diffusion-based Spatio-Temporal Graph Neural Network (DST-GNN) to the series of patient SSS networks across the three disease-stage groups. In this exploratory analysis, each sample’s COSTE-derived spatial graph was treated as a snapshot, and the model learned patterns that distinguish healthy, less affected, and more affected states. GNN explainability highlighted connections involving proliferating AT2 cells, respiratory airway secretory cells (RASC), and myofibroblasts as candidate interactions associated with the fibrotic architectural shift (Fig. 4g). Because the analysis is based on 45 samples, three stage groups, and one-hot node features, we interpret DST-GNN as a hypothesis-generating extension rather than as definitive evidence for causal drivers of disease progression. The results suggest that COSTE-derived graphs can be used as inputs for temporal modeling, while stronger validation will require larger cohorts, predefined modeling choices, and independent replication.

### 4. Nomination of spatial gene modules and niche-defined subpopulations

To further exploit COSTE’s computational efficiency, we applied it at the gene level and evaluated whether it can nominate spatially organized gene programs. We then used this capability to identify spatially restricted candidate gene programs in a human lung fibrosis dataset from systemic sclerosis patients (Fig. 5, 26 cell types annotated in a Xenium dataset)^27^. COSTE’s cell-type level analysis of one fibrotic lung sample revealed four major spatial structures, one of which corresponded to a pathologically thickened pleural region enriched for fibroblasts (cell cluster 3) and plasma cells (cluster 26) (Supplementary Fig. 12a). To identify transcripts spatially associated with this pleural niche, we performed a transcript-by-cell screening: each of the 383 detected transcript types was sequentially combined with all cell types to compute the SSS map, and its SSS was calculated relative to every cell type. We then ranked genes by their SSS to the pleural fibroblast-enriched cluster (lower SSS indicates tighter spatial co-localization within this sample). The top-ranked genes included well-known fibrosis-related factors such as IGF1 and THBS2, as well as WT1, a mesothelial transcription factor with relatively low overall abundance (Supplementary Fig. 12b, Supplementary Table 2). WT1 showed strong spatial association with the pleural structure; because WT1 is an established mesothelial marker, we interpret this as a pleural/mesothelial-associated spatial signal rather than as evidence that WT1 is a fibroblast-specific marker. The proximity of mesothelial and fibroblast-rich regions also means that segmentation or transcript-assignment effects at the pleural boundary remain possible and should be considered in follow-up validation.

**Figure 5.**
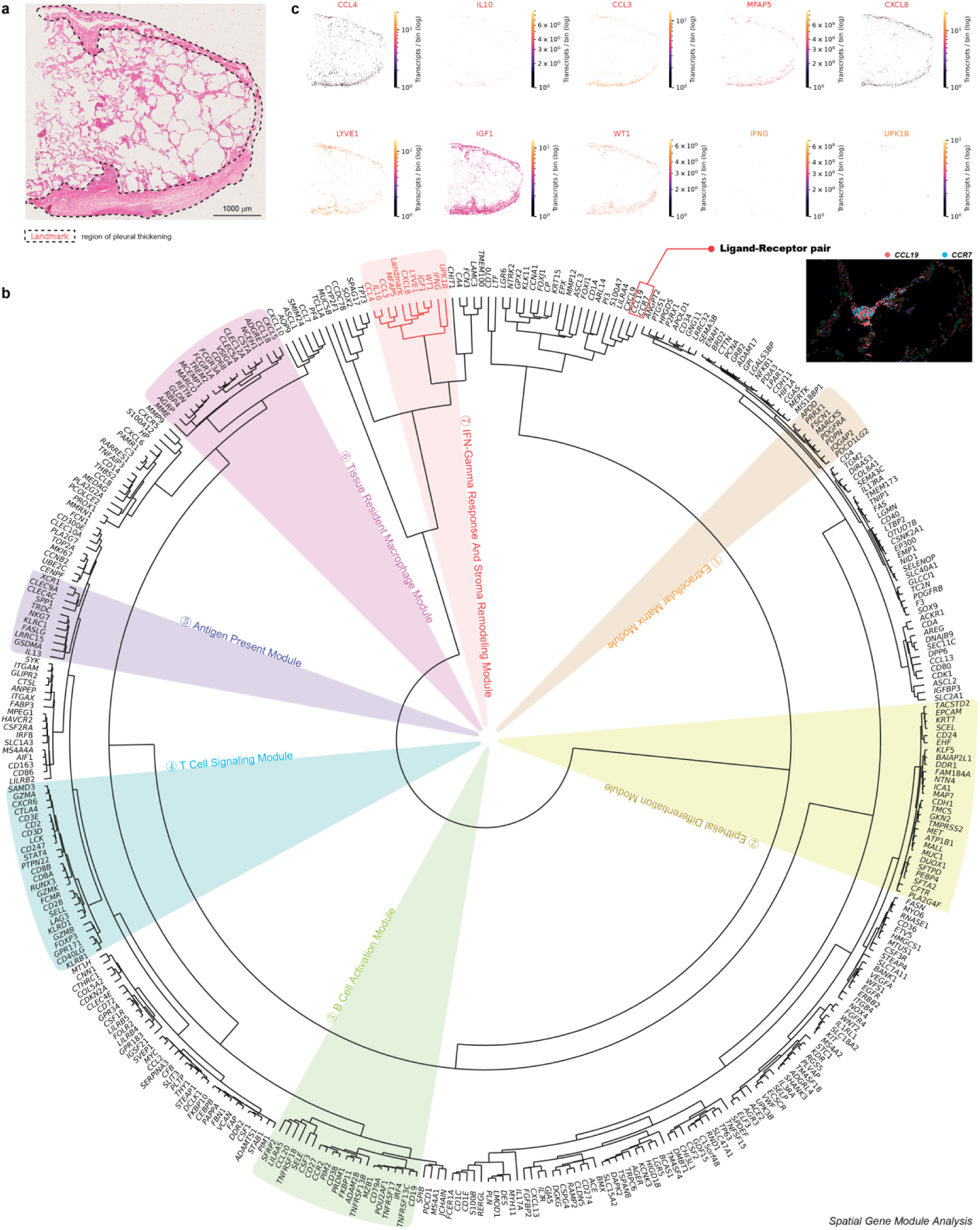
Segmentation-free COSTE analysis nominates a fibrotic pleural gene module in human lung. By incorporating tissue morphology, COSTE can operate at the single-transcript level to identify spatial gene programs without cell segmentation. (a) An H&E-stained fibrotic lung section with a thickened pleural region (dotted outline) is used as a landmark cluster for spatial analysis. (b) COSTE’s hierarchical clustering of spatial expression patterns partitions ∼400 detected transcripts into seven spatial gene modules (clusters of co-localized genes) with distinct functional themes. One module (module-7, colored red) corresponds to the fibrotic pleural niche and comprises ∼10 genes co-enriched at the mesothelial boundary, including extracellular matrix components, mesenchymal markers, and immune signaling factors. A chemokine (CCL19) and its receptor (CCR7) appear as adjacent leaves in the transcript dendrogram and are co-localized in the tissue, suggesting a spatial co-localization hypothesis for follow-up. (c) Spatial mapping of module-7 genes confirms their localization within the pleural thickening region, highlighting a pathology-associated gene program nominated by COSTE in one sample.

Among the other top spatially associated genes were IL10 and CCL4, cytokines not produced by fibroblasts but by monocytes (cluster 6). Their low SSS to the fibroblast-enriched pleural cluster indicated that IL10+CCL4+ monocytes were preferentially co-localized with the pleural fibroblast-rich region in this sample. Spatial mapping showed that monocytes immediately beneath the mesothelial layer in the thickened pleura expressed IL10 and CCL4, whereas monocytes located deeper in the lung tissue showed lower expression (Supplementary Fig. 12c). We therefore regard this pattern as a candidate pleura-associated monocyte state observed in one systemic-sclerosis lung sample. This example illustrates how transcript-level COSTE analysis can nominate cell-type-specific spatial gene programs and niche-defined candidate subpopulations, while replication in additional fibrotic samples and orthogonal validation will be needed before treating the state as a confirmed disease-associated cell population.

COSTE can also leverage histopathological context to perform morphology-guided, segmentation-free screening for lesion-associated transcripts. In the same fibrotic lung sample, we used the H&E image to identify a region of pleural fibrosis (mesothelial thickening) (Fig. 5a). We then designated all transcripts within that morphologically altered region as a single landmark cluster, without drawing any single-cell boundaries. By including this landmark as an additional cluster in COSTE’s transcript-level analysis, we probed which genes were spatially enriched in the pleural lesion. The resulting SSS dendrogram - visualized as a circular hierarchical clustering of transcripts - grouped together genes with similar spatial localizations. One module (module-7) aligned with the pleural-thickening region: in the dendrogram, the pleural landmark cluster grouped with a small set of ∼10 genes, indicating that these genes were co-localized with the pleural lesion in this sample (Fig. 5b). Spatial plotting confirmed that the module-7 genes were concentrated within the fibrotic pleura (Fig. 5c). This pleural module contained chemokines (CCL4, IL10, CCL3, CXCL8, IFNG) expressed by immune cells at the mesothelial interface^28–30^, as well as WT1 (a mesothelial cell marker^31^) and IGF1 (an epithelial and mesenchymal growth factor^32^). These co-localization patterns suggest hypotheses about inflammatory signaling at the pleural boundary. The adjacent placement of CCL19 and CCR7 in the transcript dendrogram should be interpreted as a spatial co-localization hypothesis rather than as proof of an active ligand-receptor interaction in situ. Collectively, this morphology-guided analysis demonstrates how COSTE can integrate a priori regional knowledge (*e.g.* an H&E-identified lesion) to nominate transcripts and spatial gene modules associated with a pathological region without requiring single-cell segmentation.

### 5. COSTE summarizes response-associated immune spatial topology in TNBC

Using COSTE, we analyzed imaging mass cytometry (IMC) data from the NeoTRIP TNBC trial covering baseline, on-treatment, and post-surgery samples under chemotherapy (C) or chemoimmunotherapy (C&I) arms, stratified by outcome (pCR vs RD). In this retrospective analysis, pCR cases in the chemotherapy arm showed lower SSS between PD-L1+IDO+ antigen-presenting cells (APCs) and cytotoxic CD8+GZMB+ T cells, consistent with closer relative spatial coupling of these immune populations. In the chemoimmunotherapy arm, RD tumors showed higher SSS or altered neighborhood patterns involving TCF1+ cells, Vimentin+EMT+ cells, myofibroblasts and Tregs (Supplementary Fig. 13). These treatment- and response-associated immune topologies suggest that COSTE can summarize clinically relevant spatial reorganization in the tumor microenvironment (Supplementary Figure 14). Because this analysis does not include a held-out validation cohort, leave-patient-out prediction, or ROC/AUC evaluation, we interpret these patterns as retrospective associations rather than validated response predictors.

## Discussion

We have presented a global, hierarchy-based method for analyzing tissue architecture using distance-based spatial pattern analysis. By employing a global spatial referencing strategy, COSTE summarizes tissue structure across multiple scales and at both the cell-type and transcript level. The approach can be applied to high-volume spatial omics datasets and is conceptually extensible to three-dimensional data, although 3D applications will require dedicated validation. In the datasets analyzed here, COSTE provided interpretable views of cell-cell and region-cell spatial relationships, nominated spatially confined candidate cell states, supported transcript-level analysis in unsegmented regions, and enabled morphology-guided searches for lesion-associated genes. When applied to previously published datasets, COSTE highlighted tissue structures and spatial features that complement local neighborhood-based analyses, demonstrating its utility for generating spatial biology hypotheses from existing data.

Many spatial analysis methods focus on local neighborhoods - for example, considering cells within a certain radius or a fixed number of nearest neighbors^35–39^. These approaches are well suited for short-range proximity questions, whereas COSTE is designed to provide a global summary of directed distance profiles across all annotated cell types. COSTE differs from kernel-based or Gaussian mixture approaches that compute smoothed contact frequencies because it compares distance profiles before embedding them in a hierarchical representation. This design is computationally efficient for large datasets, but it does not make the resulting scores independent of cell density, tissue size, boundary effects, or upstream cell-type annotation. Instead, COSTE should be interpreted as a complementary global representation whose results are most comparable when samples are processed with the same labels, preprocessing choices, and normalization strategy.

Despite its advantages, COSTE has several limitations that are important for interpretation. First, COSTE focuses on relative spatial pattern similarity and does not directly encode absolute distances to landmarks such as tumor boundaries, anatomical axes, or tissue edges. Such absolute measurements should be integrated as complementary features when they are biologically important. Second, SSS is a within-sample, min-max-normalized dendrogram-derived score. As a result, SSS values are most directly interpretable as relative separations within a shared analysis workflow; comparisons across samples can be informative when cell-type definitions and preprocessing are harmonized, but they should not be treated as absolute distances whose 0 and 1 values have identical meanings in every tissue. Third, the current analyses use ranking and selected group comparisons to nominate candidate spatial relationships. Formal inference will require explicit resampling frameworks, such as bootstrapping cells or fields of view to assess dendrogram and SSS stability, permutation or label-shuffle null models to test whether observed SSS deviations exceed chance expectations, and multiple-testing correction when many cell pairs or transcripts are screened. Fourth, the directed searcher-findee distance matrix is asymmetric by construction. In this manuscript, the primary SSS analyses use the searcher-based dendrogram, while the findee-based dendrogram provides a complementary perspective. Future work should systematically compare searcher-based, findee-based, and symmetrized formulations to quantify sensitivity to this choice. Fifth, nearest-neighbor distances remain sensitive to local density, rare populations, tissue boundaries and field-of-view size. Very small regions or rare compartments can yield noisy SSS patterns, and single-sample observations - including the pleural transcript-level findings - should be regarded as hypotheses until replicated in additional samples or validated by orthogonal assays.

Looking ahead, COSTE’s framework is broadly extensible across modalities and analysis contexts. We envision COSTE being used in conjunction with, rather than in replacement of, existing spatial analysis techniques. For example, one could apply COSTE to map global tissue architecture and then use higher-resolution local methods, neighborhood graphs, or gradient-based spatial gene detection to inspect particular microenvironments. The global view provided by COSTE supplies a scaffold on which to interpret local findings. The method is also applicable to multi-modal spatial data. In this study, we demonstrated its utility not only in spatial transcriptomics but also in a spatial proteomics dataset from a TNBC clinical trial24, where COSTE summarized treatment- and response-associated immune spatial reorganization. Additionally, COSTE-derived graphs can be combined with temporal modeling, as illustrated by the exploratory DST-GNN analysis, to generate hypotheses about how spatial cell-cell relationships evolve over time.

In conclusion, COSTE provides a global, hierarchy-based framework for spatial omics analysis that complements existing local approaches. By summarizing multiscale organization of cells and transcripts in tissues, COSTE provides an interpretable view of tissue architecture that can guide downstream biological investigation. Incorporating COSTE alongside neighborhood-, gradient-, and morphology-based metrics may yield a more complete understanding of tissue architecture and its alterations in health and disease. The current results support COSTE as an exploratory and hypothesis-generating tool for spatial biology, with formal statistical inference, broader benchmarking against multiscale niche methods, and independent biological validation as important next steps.

## Methods

Cophenetic Spatial Topology Embedding (COSTE) is a computational pipeline that quantifies tissue architecture by computing directed nearest-neighbor distance profiles between cell clusters and/or transcripts, constructing an inter-cluster distance matrix, and applying hierarchical clustering to summarize multilevel spatial structures. The hierarchical clustering yields an ultrametric tree (dendrogram) that encodes spatial relationships among the cell types and/or transcripts. From this tree, we derive cophenetic distances between clusters and normalize them within each analyzed sample to obtain a Spatial Separation Score (SSS) between 0 and 1. This workflow - computing directed distances, building a distance matrix, hierarchical clustering, extracting cophenetic distances, and normalizing into SSS - forms the core of COSTE. The approach does not require the user to define a spatial radius or neighborhood cutoff, and it can be scaled to large datasets using optimized nearest-neighbor search algorithms.

### Cophenetic Spatial Topology Embedding (COSTE)

We developed COSTE to embed the spatial organization of cell types and/or transcripts into a hierarchical metric space. COSTE begins by computing directed inter-type distances based on physical nearest neighbors. For each “searcher” cell of type A, we identify its nearest neighboring cell belonging to a different cell type B (the “findee”) and measure the Euclidean distance between them. By excluding same-type neighbors, this directed nearest-neighbor distance focuses on **cross-type** spatial proximity and avoids biases from cell-type self-aggregation. Formally, if *T*_*A*_ and *T*_*B*_ denote the sets of cells of type A and B, respectively, then for each cell *i* ∈ *T*_*A*_ we compute *d*(*i*, *T*_*B*_) = min_*j*∈*TB*_ dist(*i*, *j*), where dist(*i*, *j*) is the Euclidean distance between cell coordinates. Each cell of type A thus obtains a distance to type B, and averaging over all cells of type A yields a directed distance from type A to type B:

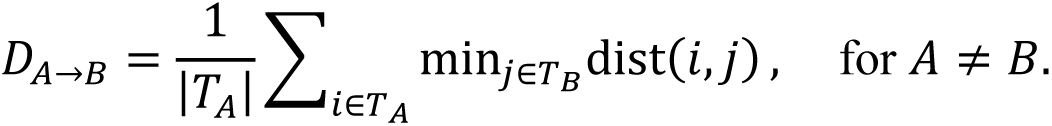

This procedure produces an asymmetric inter-cluster distance matrix *D* = [*D*_*A*→*B*_] of size *N* × *N*, where *N* is the number of cell types (matrix diagonal is undefined or set to zero by design, since no self-distances are considered). We implement the nearest-neighbor searches using efficient k-d tree based algorithms (with *k* = 1 for each target type) to ensure scalability. Instead of brute-force comparison of all cell pairs, the algorithm queries each cell type’s point set to rapidly retrieve the closest neighbor for a given cell, reducing computation time dramatically for large tissues.

Next, we use this distance matrix to reveal hierarchical structure in the spatial relationships. We apply hierarchical clustering to the matrix, treating each cell type as an observation characterized by its distance profile to all other types. Because the directed distance matrix is asymmetric, we cluster matrix rows and columns independently. Row clustering groups cell types with similar searcher profiles - that is, similar patterns of distances from that cell type to all findee types. Column clustering groups cell types with similar findee profiles - that is, similar patterns of distances from all searcher types to that target type. These two dendrograms therefore represent complementary directions of the same directed relationship rather than a single symmetric distance space. Unless stated otherwise, the main SSS heatmaps use the searcher-based dendrogram, while the findee-based dendrogram is available as a complementary output. We use average linkage (UPGMA) as the default agglomeration method. Cell types that cluster together at low linkage distances have similar directed spatial profiles, whereas those joining only at higher levels differ at broader spatial scales. This representation summarizes structure-within-structure organization without requiring a predefined neighborhood radius, but it remains dependent on the input cell-type annotations, the chosen linkage method, and the density and sampling properties of the tissue.

### Spatial Separation Score (SSS) Calculation

While the raw inter-cluster distances in the directed matrix convey which cell types are close or far from each perspective, we summarize spatial proximity using cophenetic distances from the searcher-based COSTE dendrogram. The cophenetic distance between two cell types is the linkage height at which the two types first merge in the hierarchical clustering tree. A small cophenetic distance indicates that the two cell types have similar searcher distance profiles and cluster together early, whereas a large cophenetic distance indicates that they join only at a higher level of the tree. After constructing the dendrogram, we compute pairwise cophenetic distances and normalize them to a unit interval [0, 1] within that dendrogram. Specifically, the Spatial Separation Score (SSS) is defined as:

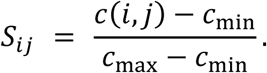

This normalization sets the lowest observed cophenetic distance in a given dendrogram to 0 and the highest observed cophenetic distance to 1. Biologically, a lower SSS indicates that two cell types have similar directed spatial profiles and are frequently co-localized or embedded within the same spatial niche in that sample. A higher SSS indicates that the two cell types are more separated in the sample-specific hierarchy or occupy distinct tissue compartments. Because the normalization is performed within each dendrogram, SSS should be interpreted as a relative, sample-normalized score. Cross-sample comparisons are most meaningful when the same cell-type definitions, preprocessing, and analysis workflow are used, and even then they should be interpreted as relative architectural shifts rather than absolute physical distances. We assemble pairwise SSS values into an SSS matrix and visualize it as an SSS heatmap to summarize global spatial organization. Clusters of cell types that form integrated spatial communities appear as blocks of low SSS, whereas more segregated cell-type pairs show high SSS values. All downstream analyses of spatial patterns, including identification of nested structures and distinct microenvironments, are based on this sample-normalized SSS matrix.

### Diffusion-based Spatio-Temporal Graph Neural Network (DST-GNN) Analysis

Graph neural networks (GNNs) naturally capture graph-structured relationships and have been applied to spatial omics analysis^40, 41^. In this study, we used a Diffusion-based Spatio-Temporal Graph Neural Network (DST-GNN) as an exploratory extension for graph-based spatio-temporal analysis. We compiled spatial relationship data from a cohort of 45 tissue samples, grouped into three disease-stage categories (T1: healthy; T2: less affected; T3: more affected). Each sample’s spatial architecture was represented as a graph in which the 47 predefined cell types served as nodes, with the SSS between each pair of cell types used as the edge weight (ranging from 0 to 1, with lower values indicating closer relative spatial proximity within the sample). If a given pair of cell types had no notable spatial association in a sample, we treated the edge between them as absent by assigning it a weight of 1 for that sample.

Using this collection of weighted graphs across disease-stage groups, we trained a DST-GNN model to capture patterns in cell-cell spatial relationships. In our implementation (built with PyTorch Geometric), each cell type is represented as a node with a one-hot encoded feature vector (a 47-dimensional identity vector since each node’s type is unique). The model employs graph convolutional layers with diffusion-based message passing, which allow information to propagate through the graph in a way that accounts for edge weights and graph structure. We organized the data as a temporal sequence of graph groups (T1 to T2 to T3) and optimized the model with Adam to minimize a mean squared error loss when predicting or reconstructing later-stage spatial connection patterns from earlier-stage patterns. Given the modest sample size, limited number of stage groups, and simple node features, this model was used for exploratory prioritization rather than formal prediction.

After training the DST-GNN, we applied GNNExplainer to identify nodes and edges that were influential for the model’s reconstruction of stage-associated graph changes. This analysis highlighted candidate cell types and interactions whose SSS-derived edge weights changed across T1, T2 and T3. We visualized these candidate interactions to facilitate biological interpretation, but we do not treat them as statistically validated drivers of tissue remodeling. Larger cohorts, explicit train/validation/test splits or cross-validation, baseline temporal models, and independent replication will be required before DST-GNN results can be used for formal inference about disease progression.

### Benchmark dataset simulation

We simulated a collection of synthetic 2D spatial datasets to serve as ground-truth benchmarks for spatial association analysis. All simulations were conducted on a normalized square plane (unit square coordinates) populated by six distinct cell types (labeled A through F). Each dataset was generated either with equal numbers of each cell type or with specified proportions, depending on the scenario, and multiple realizations were produced using different random seeds to ensure robust testing of method performance. We designed three qualitatively different spatial patterns – **modular**, **nested**, and **random** – to represent, respectively, strongly co-localized niches, hierarchical concentric layering, and a completely unstructured spatial distribution.

### Modular pattern

The modular configuration consisted of three well-separated cell clusters (“modules”) arranged in space, each module containing a specific pair of cell types co-mingled. In a base configuration, cell types A and B formed Module 1, types C and D formed Module 2, and types E and F formed Module 3. The module centers were positioned far apart relative to their cluster size (for example, placing Module 1 at the origin (0,0), Module 2 at a distant coordinate along the x-axis, and Module 3 offset in the y-direction), so that each module was largely isolated from the others. Within each module, cells of the two corresponding types were placed in close proximity by sampling their coordinates around the module’s center point – for instance, by drawing displacements from a bivariate normal distribution centered at the module centroid (with a chosen standard deviation controlling the cluster’s radius). This procedure produces a tight clustering of each cell-type pair, effectively creating three distinct co-localized niches (A–B, C–D, and E–F) with minimal intermixing between modules. To explore variations of this pattern, we perturbed key parameters and generated alternative modular datasets. In one set of variants, we varied the distance between modules (e.g., decreasing the separation to bring clusters closer or increasing it to push them further apart) to assess the effect of module proximity on analysis outcomes. In another variant, we altered the pairing scheme of cell types – for example, swapping partner types between modules – thereby testing scenarios with different co-localization pairings. We also modified the composition of certain modules in some simulations, such as adjusting the relative abundance of the two cell types within a module or introducing a small fraction of an “out-of-place” cell type into a module, to examine the robustness of methods to impurities or uneven cell-type ratios. All these modular pattern datasets share the defining feature of discrete, localized cell-type pairs, providing controlled cases where the expected true associations (within-module pairs) are known by construction.

### Nested pattern

In contrast to the modular clusters, the nested pattern was designed to mimic a hierarchical, concentric organization of cell types (akin to layered tissue zones). Each nested dataset contained two independent multi-layer structures, defined as triplets of cell types A–B–C and D–E–F, respectively. Within each triplet, the three cell types were arranged as concentric shells around a common center: one cell type occupied a small inner core region, a second cell type formed a middle ring encircling that core, and a third cell type formed an outer ring around the middle layer. For example, in one triplet, type A might occupy the center, type B the middle annulus, and type C the outer annulus (denoted A ⊂ B ⊂ C, indicating that A is enclosed by B, which is enclosed by C). The second triplet was constructed similarly (D ⊂ E ⊂ F) and placed at a different location in the plane. We defined each shell in polar coordinates as an annulus with specified inner and outer radii from the triplet’s center. Cell coordinates for a given shell were then sampled uniformly at random within that annular region by drawing a radius *r* from a uniform distribution between the shell’s inner and outer radius and an angle θ from a uniform distribution on [0, 2π), and converting *(r, θ)* to Cartesian *(x, y)*. This sampling scheme yields an even distribution of points throughout each shell and ensures a clear, sharp boundary between adjacent layers. All points belonging to the same shell were assigned the corresponding cell-type label (A–F), so that the ground-truth spatial relationships among cell types are fully determined by this layered construction (e.g., A is physically proximal to B, and B to C, by virtue of nesting, whereas A and C are farther apart, being separated by an intervening layer). Unless noted otherwise, we kept the number of cells per type consistent across comparable nested simulations by assigning, for example, a smaller number of points to the core types (A and D), a moderate number to the middle-layer types (B and E), and a larger number to the outer-layer types (C and F) – reflecting the increasing area of concentric shells – while ensuring that each corresponding pair of shells (A vs D, B vs E, C vs F) had equal cell counts. This yielded six cell types with either equal or proportional population sizes (depending on the scenario) and maintained symmetry between the two triplets A–B–C and D–E–F so that they were structurally identical apart from their spatial placement.

We introduced several controlled perturbations to the nested pattern to create a family of variant datasets and to test method sensitivity to different geometric features. First, in the **shell-spacing variants**, we altered the radial spacing between the inner, middle, and outer shells. In some simulations the shells were tightly packed (small gaps between the concentric layers), whereas in others the shells were pushed farther apart by increasing the difference between the inner and outer radii of consecutive layers. This resulted in nested structures ranging from very compact (all three layers in close proximity) to very expanded (large radial distance separating the core, middle, and outer rings). Second, in the **shell-thickness variants**, we varied the thickness of the rings comprising the middle and outer layers. Here, one extreme condition had very thin shells (each layer occupying only a narrow radial band), while the opposite extreme used much thicker shells (each layer spanning a wide radial range), with intermediate cases in between. By modulating annulus thickness, we effectively changed the density and spatial extent of each layer, allowing us to examine whether analysis methods are affected by how broad or narrow a spatial domain a given cell type occupies. Third, in the **inter-group distance variants**, we manipulated the distance between the two triplet structures (A–B–C vs. D–E–F). In the closest configuration, the two nested triplets were placed relatively near each other in the field of view (even partly overlapping in their outer neighborhoods), whereas in the most distant configuration, they were located far apart (well-separated such that each triplet formed its own distinct region in the plane). Intermediate placements were also considered, producing a graded series from minimal separation to large separation between the two groups. Varying the inter-group distance tests whether a method can distinguish two separate but similar structures and whether it erroneously merges them or correctly treats them as independent depending on proximity. Across all these nested pattern simulations, the fundamental “ground truth” remains that A, B, C form one hierarchical cluster of concentric layers and D, E, F form a second, analogous cluster. Thus, a successful spatial association analysis should detect strong associations (tight spatial coupling) among the cell types within each triplet (e.g., A–B, B–C, and indirectly A–C through the hierarchy) and little to no association between the two different triplets when they are well separated. The nested pattern series provides a rigorous assessment of whether algorithms can recover layered spatial architectures and remain stable under changes in scale (shell radii), layer thickness, and between-group spacing.

Random pattern: In addition to the structured modular and nested scenarios, we generated a random pattern dataset in which all six cell types A-F were uniformly randomly distributed over the unit square with no underlying spatial structure. For this null model, each cell’s (x, y) coordinates were sampled independently from a uniform distribution on the [0,1] x [0,1] field, and cell types were assigned in equal proportion (or according to a specified global frequency, if applicable) but without any spatial bias - meaning that the location of a cell was independent of its type. This random configuration produces a well-mixed tissue with no true cell-cell preferential arrangements or gradients. It serves as a baseline to confirm that spatial association methods do not spuriously detect structure where none exists: ideally, analysis of the random pattern should yield no preferential associations between particular cell-type pairs beyond what is expected by chance.

In summary, our synthetic dataset collection spans three fundamental spatial organization regimes – discrete co-localized clusters, concentric layered domains, and fully random mixing. By constructing these patterns in a mathematically controlled manner and introducing systematic perturbations (in inter-module distance, shell radii, layer width, and inter-group positioning), we created an array of test cases with known ground-truth relationships. This allows us to benchmark spatial association analysis tools by evaluating whether they can correctly identify the intended cell-type associations (e.g., the paired modules or nested shells) and how robust their performance remains under changes in spatial scale, cell density distribution, or the degree of separation between structures. The use of multiple random seeds for each configuration further ensures that results are not driven by one particular point arrangement but are representative of the underlying pattern. These synthetic benchmarks thus provide a solid validation framework for comparing methods and assessing their sensitivity and specificity in detecting spatial associations under various realistic scenarios.

### Benchmarking of spatial structure detection on synthetic datasets

We compared four approaches - Squidpy NE, Giotto NE, ANE, and COSTE - on synthetic spatial datasets with known modular and nested structures. Squidpy, Giotto and ANE were used here as local neighborhood-enrichment baselines, because they summarize cell-type co-occurrence within local neighborhoods. Each method was given the same input consisting of the cells’ spatial coordinates and their ground-truth cell type labels. Squidpy and Giotto use permutation-based neighborhood enrichment tests (Squidpy’s default 1,000 permutations; Giotto with 1,000 permutations in our tests) to calculate enrichment statistics, whereas ANE computes enrichment via an exact analytical formula. COSTE does not require specifying a neighborhood radius or neighbor count for the core SSS calculation; instead, it produces a sample-normalized SSS matrix from directed distance profiles and hierarchical clustering. We generated neighborhood enrichment matrices and COSTE SSS matrices to compare how local enrichment summaries and global hierarchical summaries represent modular, nested and random synthetic geometries. This benchmark is intended to characterize behavior relative to local NE baselines, not to replace head-to-head comparisons with methods specifically designed for hierarchical or multiscale niche discovery.

### Runtime and memory benchmarking on a real tissue dataset

We next evaluated the computational efficiency of each method on a large, real-world tissue dataset: the Xenium neonatal mouse pup, which contains 1,296,241 cells encompassing 44 distinct cell types. All four methods were applied to the full dataset using the same inputs (the 2D coordinates of each cell and its assigned cell type label, provided as an AnnData object). For Giotto’s neighbor enrichment, we set number_of_simulations = 100 (i.e., 100 permutation runs for significance) and used a k = 10 nearest-neighbor definition to mirror a typical neighborhood size. Squidpy’s spatial enrichment was run with k = 10 and 1,000 permutations. The ANE method was applied in its analytical mode with no permutations required, and COSTE was executed using its standard analytical pipeline. Each method produces a cell type x cell type matrix capturing spatial associations among cell types - for example, Squidpy and Giotto report enrichment as standardized Z-score matrices or significance values for each pair, while COSTE outputs the SSS matrix of pairwise relative separation scores.

### Squidpy neighborhood enrichment analysis with varying parameters

In Squidpy NE, spatial proximity is assessed by examining each cell’s immediate neighborhood rather than using global distance profiles. We constructed cell–cell neighbor graphs for the mouse pup dataset and evaluated multiple neighborhood definitions to test the impact of parameter choices. Specifically, cells were connected either to their *k* nearest neighbors (*k* = 4, 6, 8, 10, or 15) or to all neighboring cells within a fixed radial distance (50 µm or 100 µm). For each choice of *k* or radius, we then computed an enrichment score for every pair of cell types by comparing the observed frequency of their neighbor interactions to a random expectation obtained by permutation testing. Neighborhood enrichment scores for a given cell-type pair were derived by contrasting the observed neighbor counts with this null distribution.

## Author Contributions

M.L., T.H., and M.N. conceived and supervised the study. T.H. and A.S. performed the data analysis. M.L., T.H., and M.N. wrote the manuscript. A.S. and C.S. contributed to data analysis, reviewed the manuscript, and provided critical feedback.

## Acknowledgements

This work was supported by grants to M.N. from Horizon Europe Mission on Cancer “SPACETIME” project (grant no 101136552), Cancerfonden (24-3457), the Swedish Research Council (2024-02533), U-CAN, and the CW R&D fund. A.S. and C.S. were supported by Cancerfonden 24-3852. The computations were enabled by resources provided by the National Academic Infrastructure for Supercomputing in Sweden (NAISS), partially funded by the Swedish Research Council through grant agreement no. 2022-06725.

## Data Availability

All spatial transcriptomic datasets used in this study are publicly available. The Xenium Prime FFPE neonatal mouse pup dataset is available from 10x Genomics at https://www.10xgenomics.com/datasets/xenium-prime-ffpe-neonatal-mouse, and the Xenium Prime fresh-frozen mouse brain coronal section dataset at https://www.10xgenomics.com/datasets/xenium-prime-fresh-frozen-mouse-brain. The Xenium lung fibrosis dataset used in Figures 1 and 2 is available in the Gene Expression Omnibus (GEO) under accession number GSE276945, and the lung fibrosis sample from a systemic sclerosis patient used in Figure 3 under accession number GSE303048 (patient “Ssc2”).

## Code Availability

The method presented in this manuscript is COSTE. The primary Python software distribution is released as Cell-GPS, and the corresponding Python import package is cellgps, available at: https://github.com/hutaobo/cellgps. This repository contains the Python implementation, documentation and example workflows used for the main analyses. A lightweight R implementation for R users is provided separately at: https://github.com/hutaobo/cellgpsr; it implements core COSTE functionality but does not expose the full Python feature set. In addition, a compiled Windows executable with a graphical user interface is archived on Zenodo (https://zenodo.org/records/17859173). For reproducibility, users should cite the specific repository release or Zenodo record used for analysis.

## Ethics Statement

This study did not involve any new experiments with human participants or live animals. All analyses were performed on datasets from previously published studies, each of which obtained informed consent and ethical approval as needed. No additional ethical approval was required for the purely computational work presented here.

## Supplementary Information

Supplementary Information is provided with this submission, including additional figures and tables that support the findings of this study.

## Supplementary Figures

**Supplementary Figure 1.**
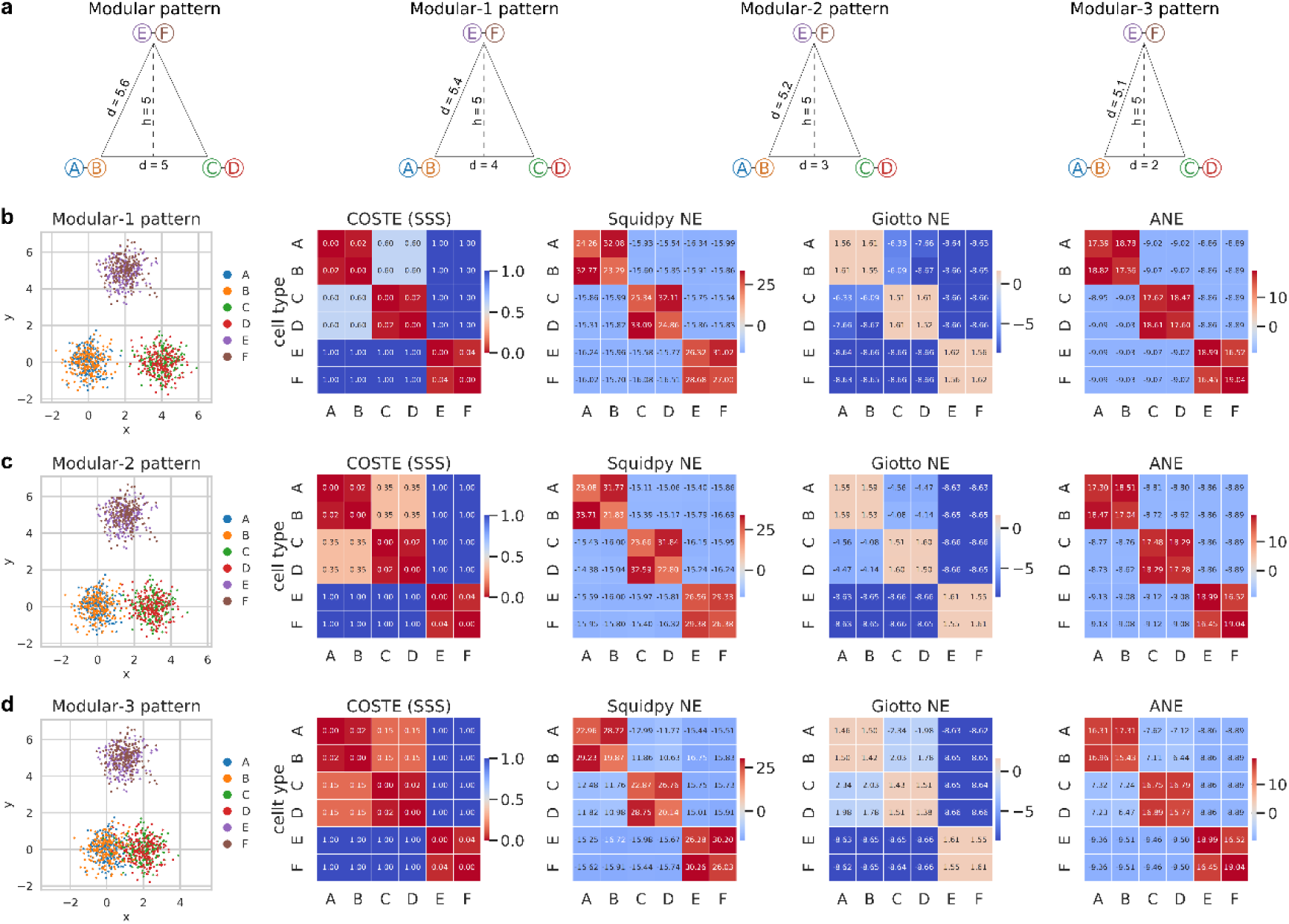
Synthetic modular spatial patterns and benchmarking results. Synthetic 2D modular datasets in which six cell types (A-F) form three well-separated co-localized modules are used to compare COSTE with local neighborhood-enrichment baselines. The panels show the simulated spatial layouts together with the corresponding cell-cell association heatmaps for COSTE, Squidpy NE, Giotto NE and ANE; all four approaches recover the expected block-diagonal modular structure.

**Supplementary Figure 2.**
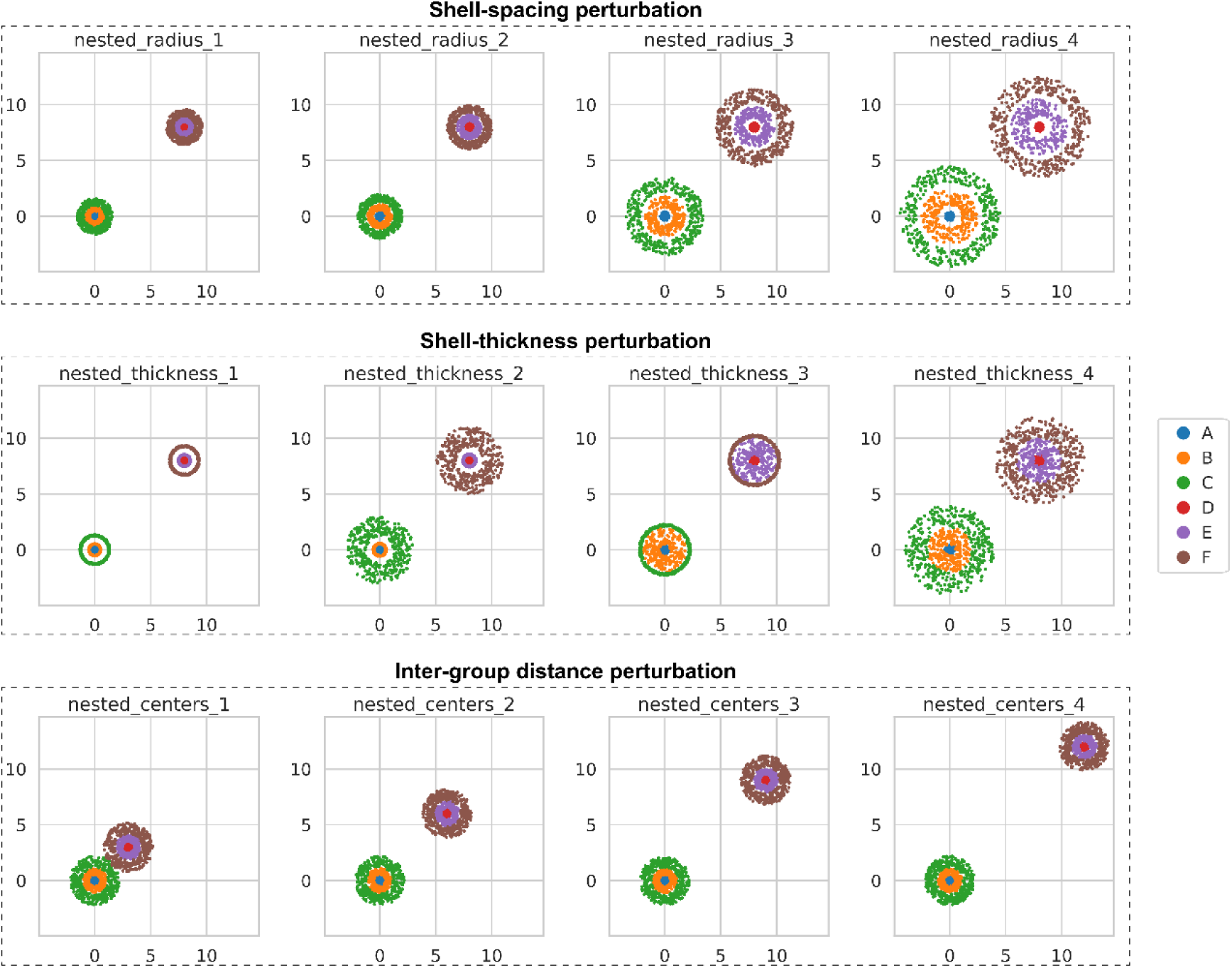
Synthetic nested spatial patterns used for benchmarking spatial association methods. This figure illustrates the synthetic “nested” spatial patterns used to benchmark COSTE against neighborhood-based spatial association methods. Each dataset contains two independent triplet structures, A–B–C and D–E–F, which represent multi-layered spatial domains. Within each triplet, three concentric shells form an inner core, a middle ring and an outer ring around a common center. Points are sampled uniformly at random within each annulus by drawing radii from a uniform distribution between the specified inner and outer radii and angles from a uniform distribution on [0, 2π), and then converting to Cartesian coordinates. All points in a given shell are assigned the same cluster label, resulting in six cell types (A–F) with well-defined ground-truth spatial relationships. To probe how spatial methods respond to changes in global scale, shell thickness and inter-structure separation, we generated twelve nested configurations grouped into three perturbation families. In the shell-spacing series (nested_radius_1–4), we progressively increased the radii of the middle and outer shells while keeping the inner core fixed, producing patterns that range from compact to widely separated nested structures. For example, nested_radius_1 uses radial intervals of (0–0.4), (0.45–0.9) and (0.95–1.6) for the core, middle and outer shells, respectively, whereas nested_radius_4 expands these intervals to (0–0.4), (1.0–2.5) and (3.0–4.5). In the shell-thickness series (nested_thickness_1–4), we altered the widths of the middle and outer annuli to generate extremely thin or thick layers; nested_thickness_1 employs narrow shells with radii (0.5–0.55) and (1.2–1.35), while nested_thickness_4 uses much thicker shells with radii (0.5–2.0) and (2.0–4.0). In the inter-group distance series (nested_centers_1–4), the two triplets are placed at increasing distances in the plane, with centers at (0, 0) versus (3, 3), (6, 6), (9, 9) and (12, 12), thereby creating scenarios where the two nested structures range from partially overlapping neighborhoods to clearly separated domains. Together, these synthetic patterns provide a controlled and interpretable benchmark for evaluating whether spatial association methods correctly recover the nested A–B–C and D–E–F architectures, remain stable under changes in shell spacing and thickness, and appropriately distinguish or merge the two triplets as their separation varies. COSTE, Squidpy neighborhood enrichment, Giotto proximity enrichment and ANE are all applied to this shared set of geometries in subsequent benchmarking analyses.

**Supplementary Figure 3.**
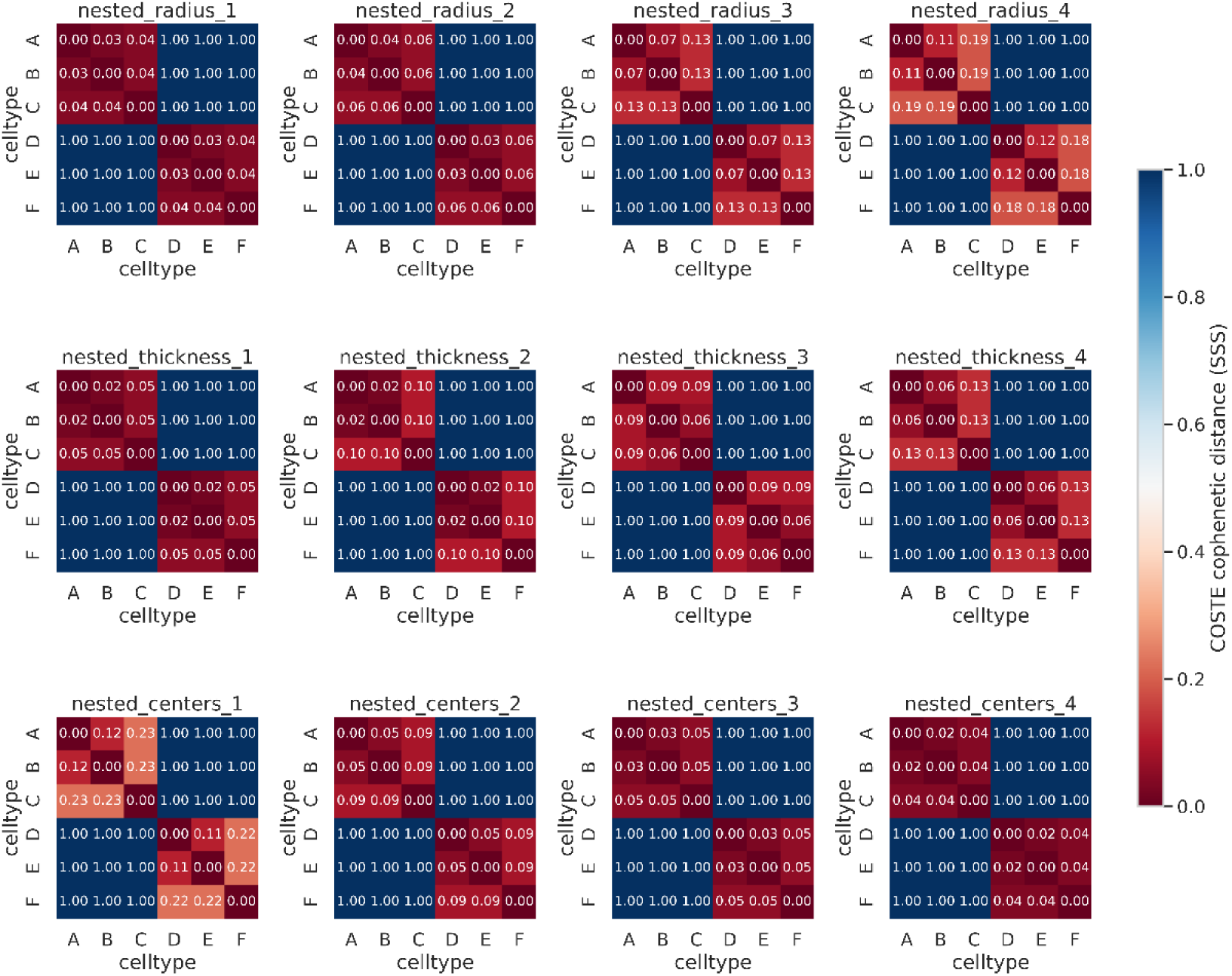
COSTE analysis of synthetic nested spatial patterns. Spatial Separation Score (SSS) heatmaps produced by COSTE for the twelve nested configurations described in Supplementary Fig. 2 (shell-spacing, shell-thickness and inter-group distance variants). Across these variants, COSTE preserves the two triplets of nested layers (A-B-C and D-E-F) as distinct structures in the SSS representation under the tested simulation settings.

**Supplementary Figure 4.**
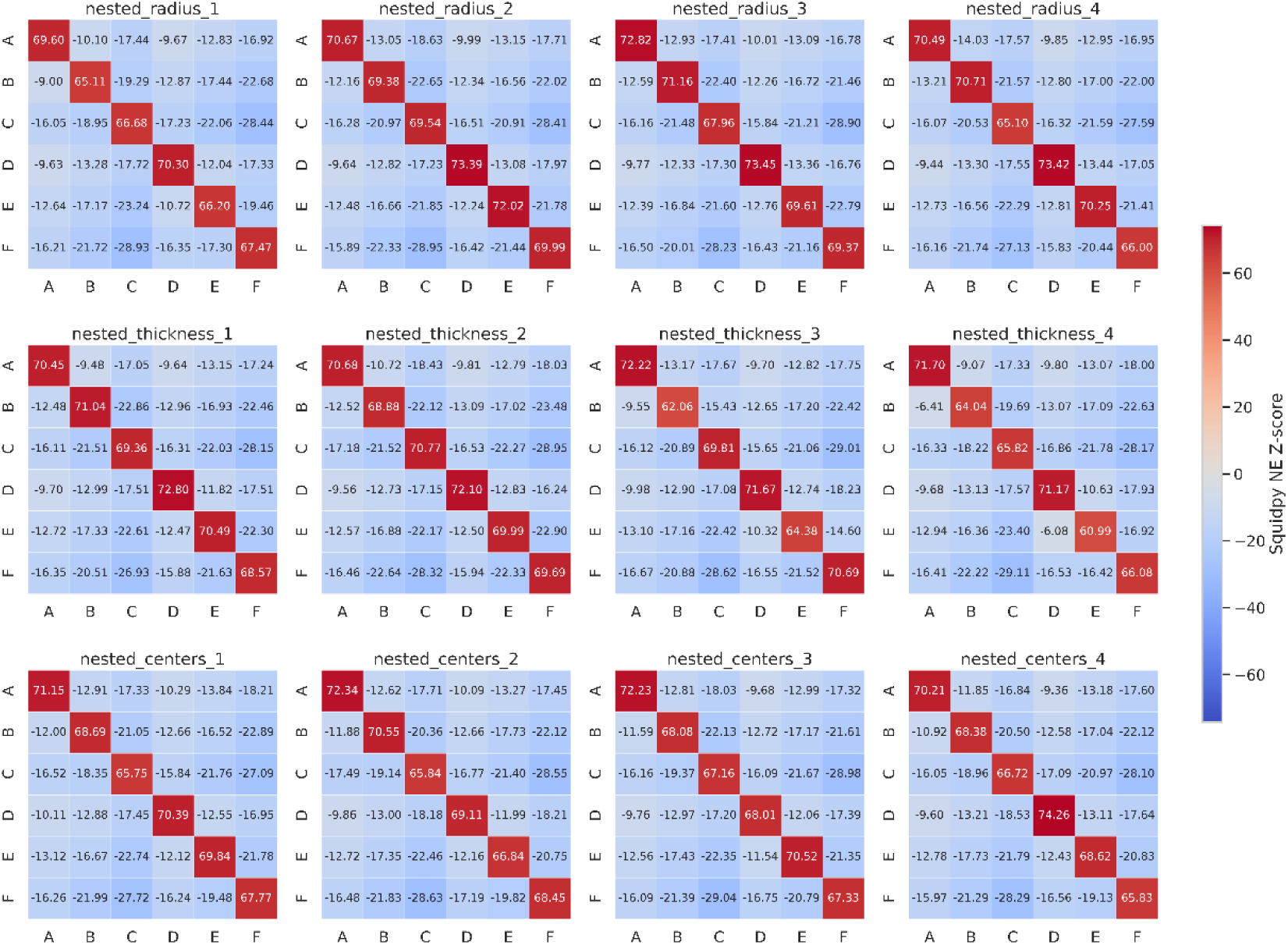
Squidpy neighborhood-enrichment analysis of synthetic nested spatial patterns. Neighborhood-enrichment heatmaps generated by Squidpy NE for the twelve nested configurations in Supplementary Fig. 2. Across these variants, Squidpy does not recover the two nested triplets (A-B-C and D-E-F) as separate spatial modules under the tested settings, illustrating how a local neighborhood summary can be less suited to this specific hierarchical nested geometry.

**Supplementary Figure 5.**
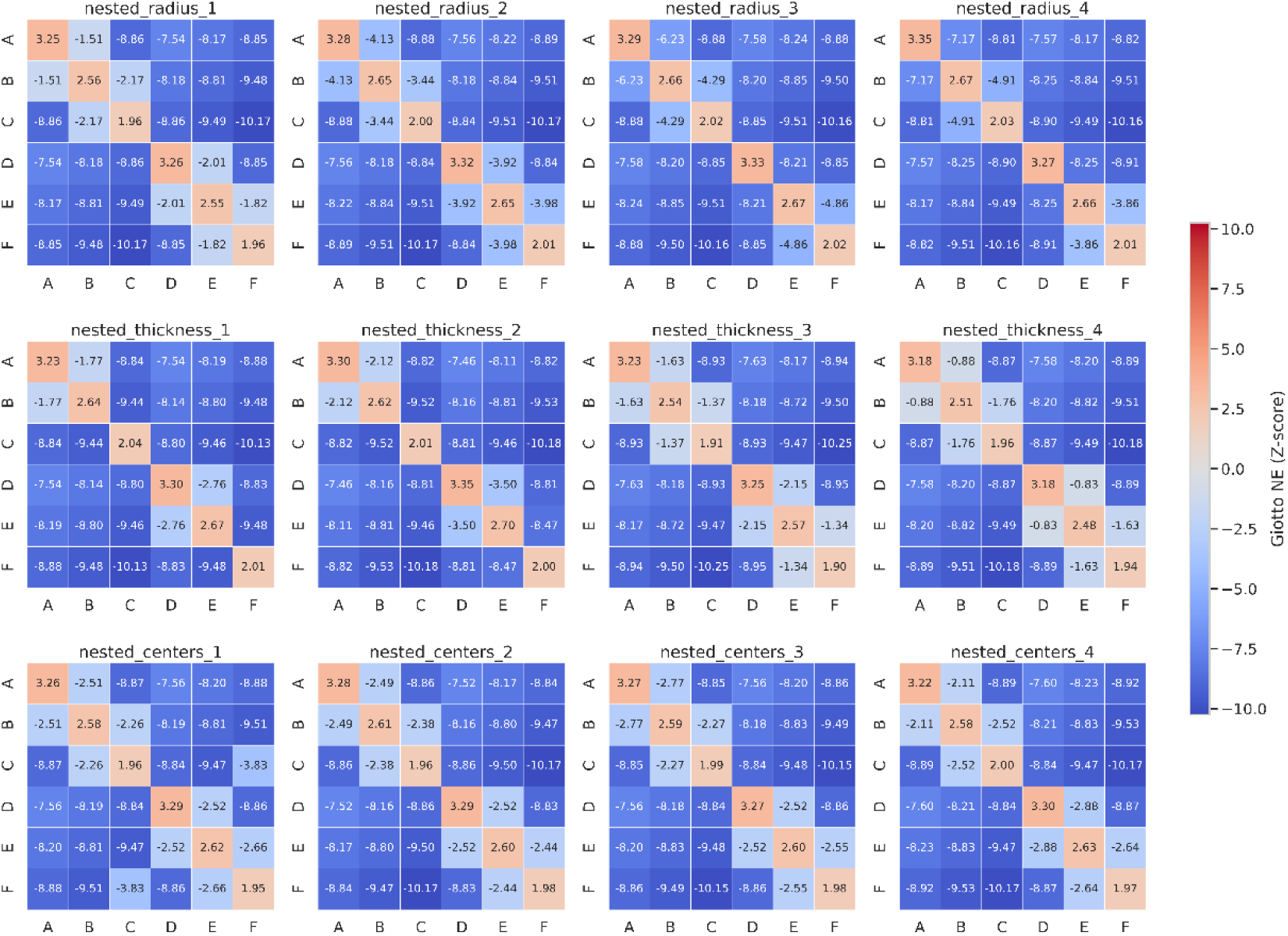
Giotto neighborhood-enrichment analysis of synthetic nested spatial patterns. Neighborhood-enrichment heatmaps generated by Giotto NE for the twelve nested configurations in Supplementary Fig. 2. Giotto occasionally identifies enriched pairs of cell types, but under the tested settings it does not consistently delineate the two nested A-B-C and D-E-F structures as complete modules.

**Supplementary Figure 6.**
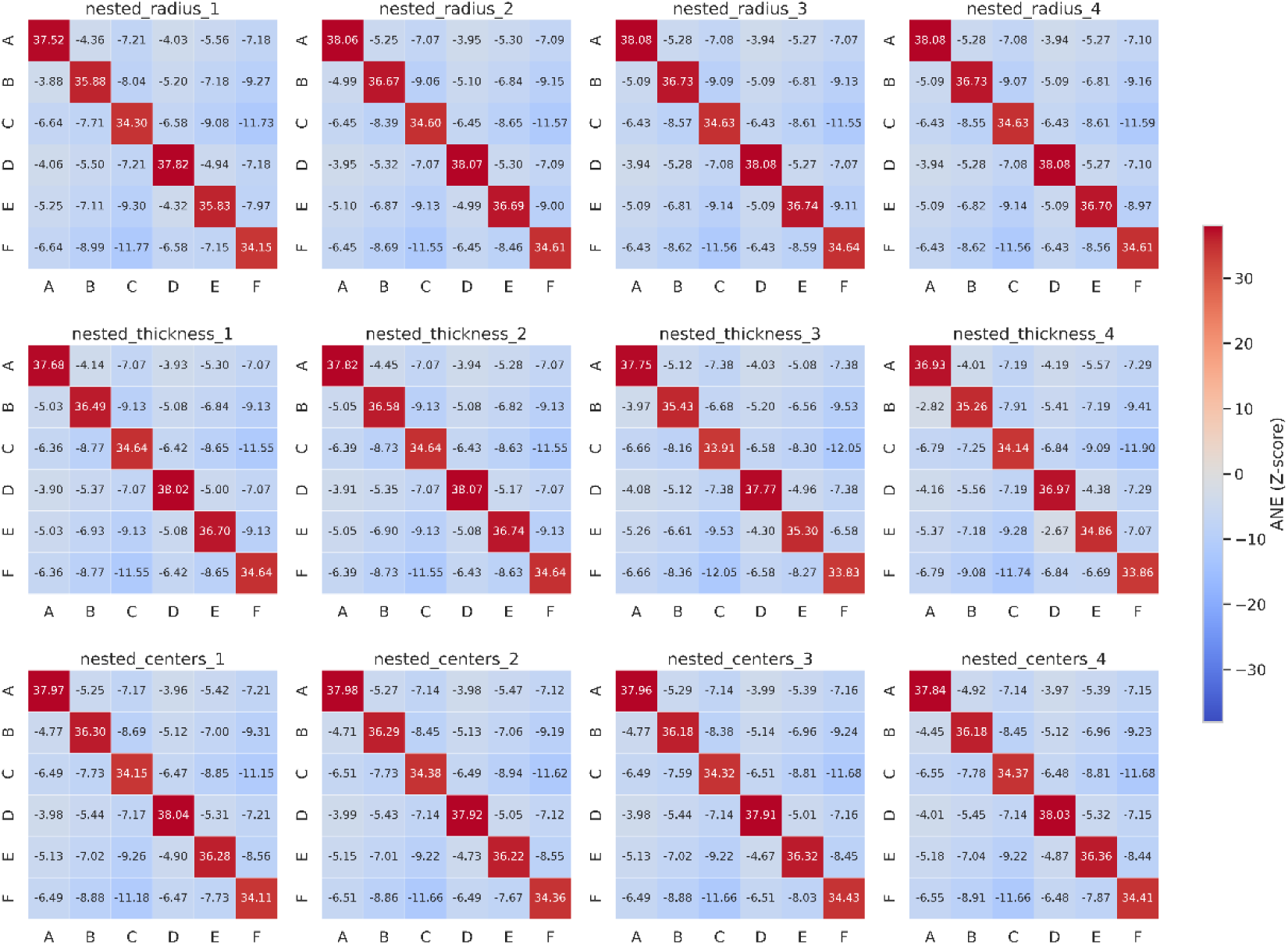
ANE analysis of synthetic nested spatial patterns. Analytical Neighborhood Enrichment (ANE) association matrices for the twelve nested configurations in Supplementary Fig. 2. Under the tested settings, ANE does not consistently separate the two nested triplets (A-B-C and D-E-F) into distinct spatial modules, whereas COSTE preserves the underlying nested structure in its SSS representation.

**Supplementary Figure 7.**
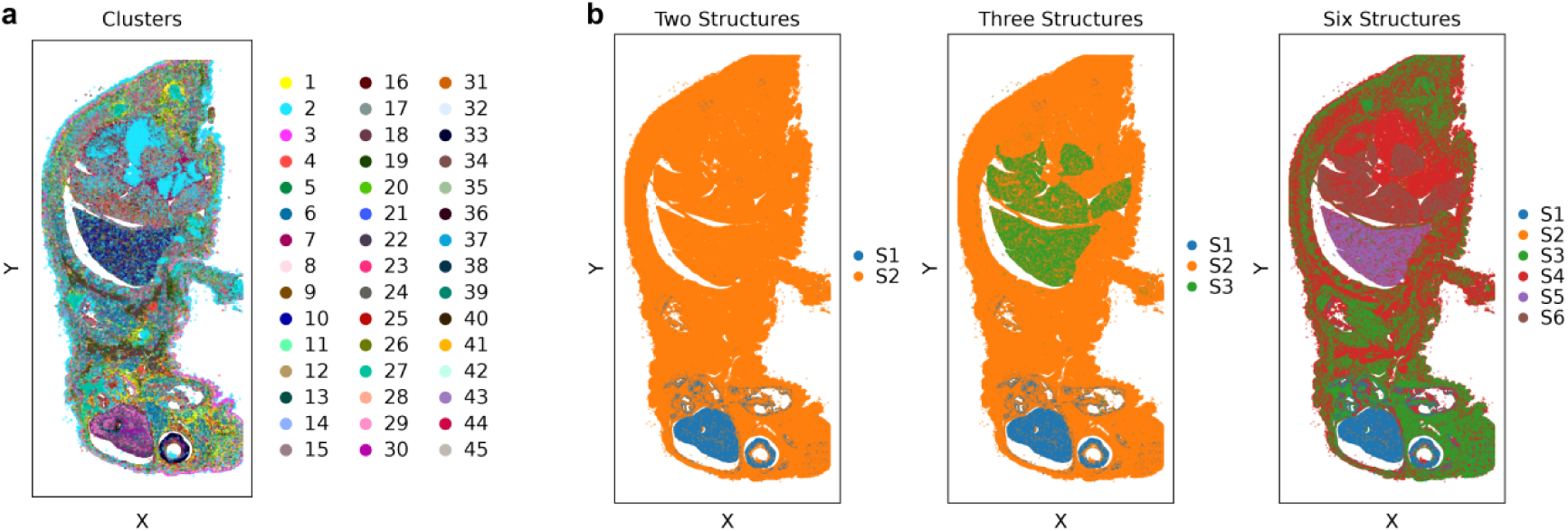
Hierarchical spatial structures identified by COSTE in a neonatal mouse pup. (a) All 45 cell types profiled in the neonatal mouse tissue are mapped onto the histological section by their transcriptomic identities, with each cell cluster labeled and color-coded. This provides a global view of the tissue’s cellular organization. (b) Spatial structures inferred by COSTE at increasing levels of resolution: two broad structures (S1, S2), three structures (S1-S3), and six finer structures (S1-S6). These partitions reveal a structure-within-structure organization: as the number of identified structures increases, smaller and more cohesive domains emerge within larger regions. This multiscale hierarchy illustrates COSTE’s ability to summarize tissue architecture across multiple spatial scales without requiring a predefined neighborhood radius.

**Supplementary Figure 8.**
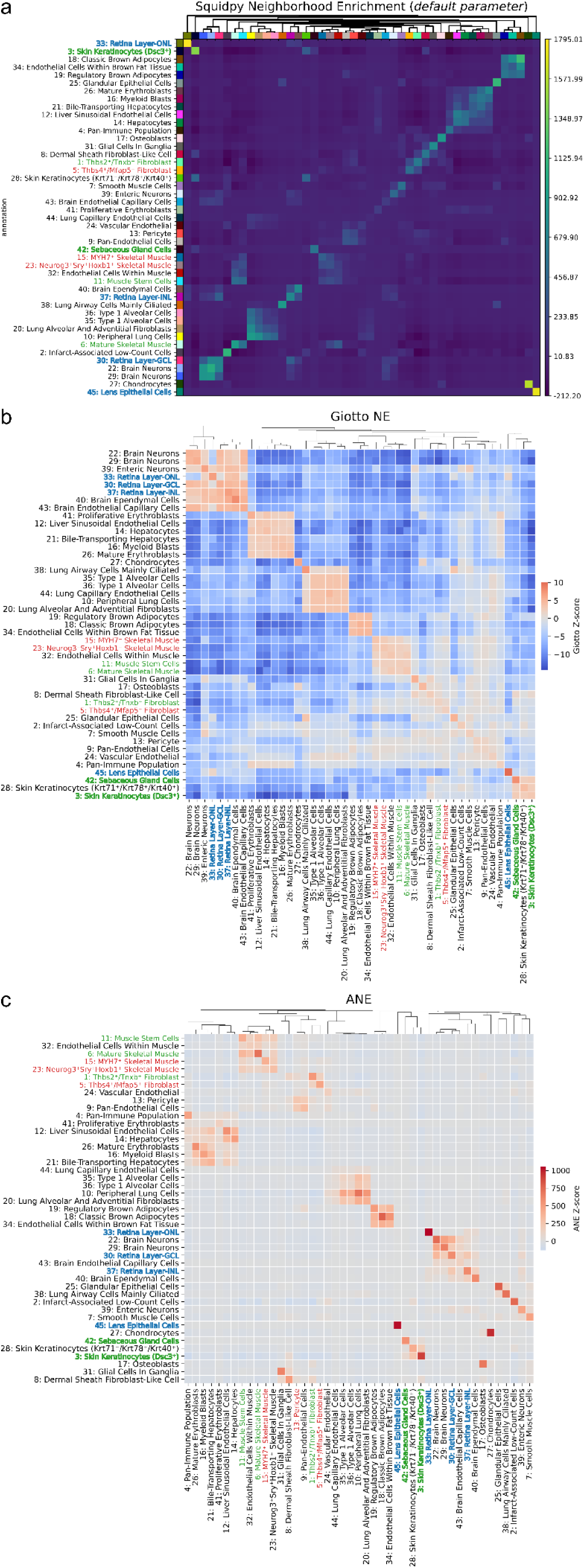
Comparing Squidpy, Giotto and ANE neighborhood-enrichment methods in a neonatal mouse pup. For the neonatal mouse Xenium dataset, neighborhood-enrichment heatmaps from Squidpy NE (a), Giotto NE (b) and ANE (c) provide local cell-type association summaries that differ from the global COSTE hierarchy. Cell types that are co-localized in the tissue map can be separated in the clustered neighborhood-enrichment heatmaps, indicating that local and global spatial representations capture different aspects of the data.

**Supplementary Figure 9.**
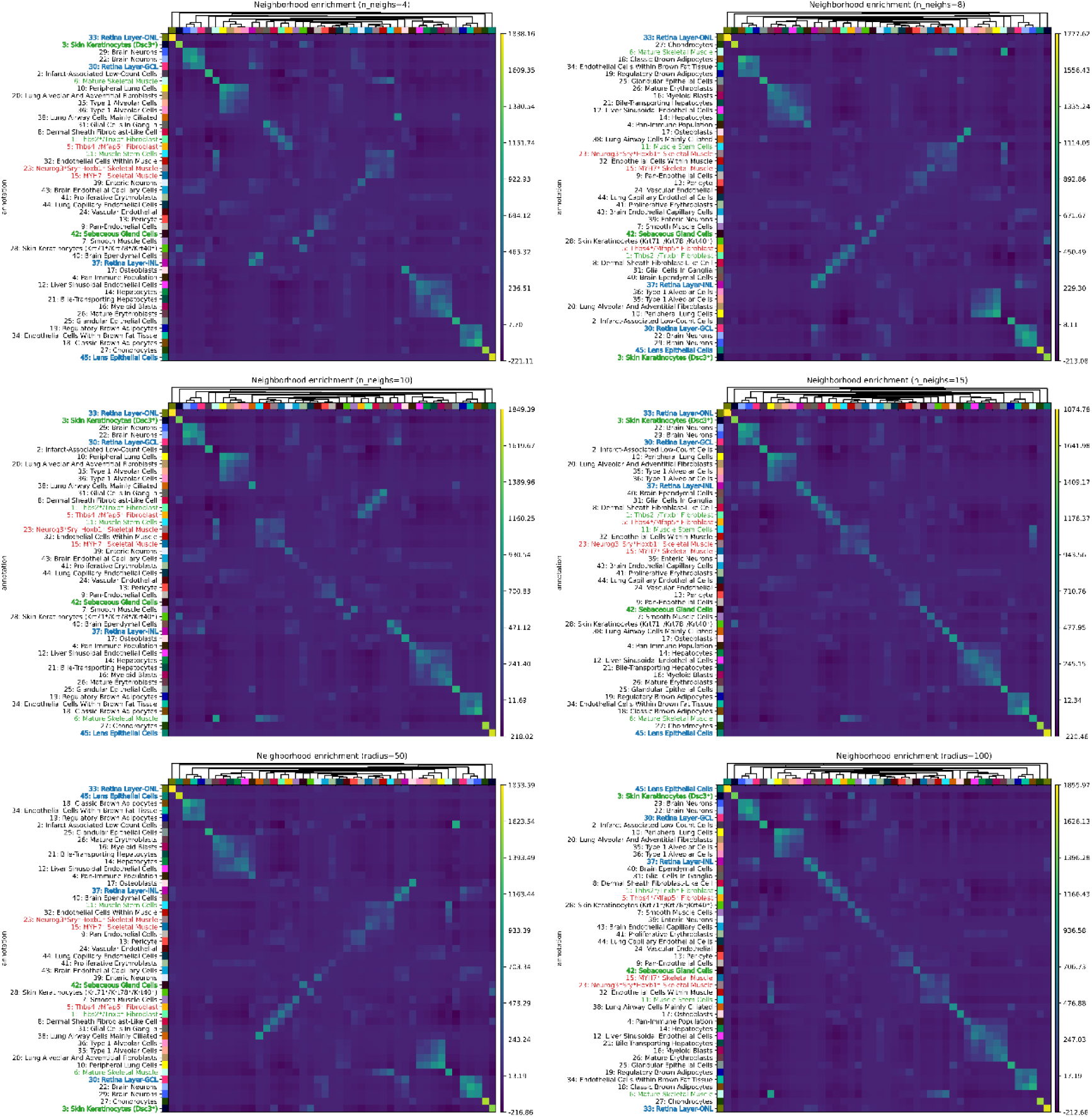
Squidpy neighborhood-enrichment analysis with varied parameters in a neonatal mouse pup. For the neonatal mouse Xenium dataset, Squidpy neighborhood-enrichment heatmaps are shown under multiple parameter settings (varying the number of nearest neighbors and the neighborhood radius). Across the tested parameters, Squidpy shows strong self-enrichment along the diagonal and parameter-sensitive off-diagonal association patterns, highlighting the dependence of local neighborhood summaries on the chosen neighborhood definition.

**Supplementary Figure 10.**
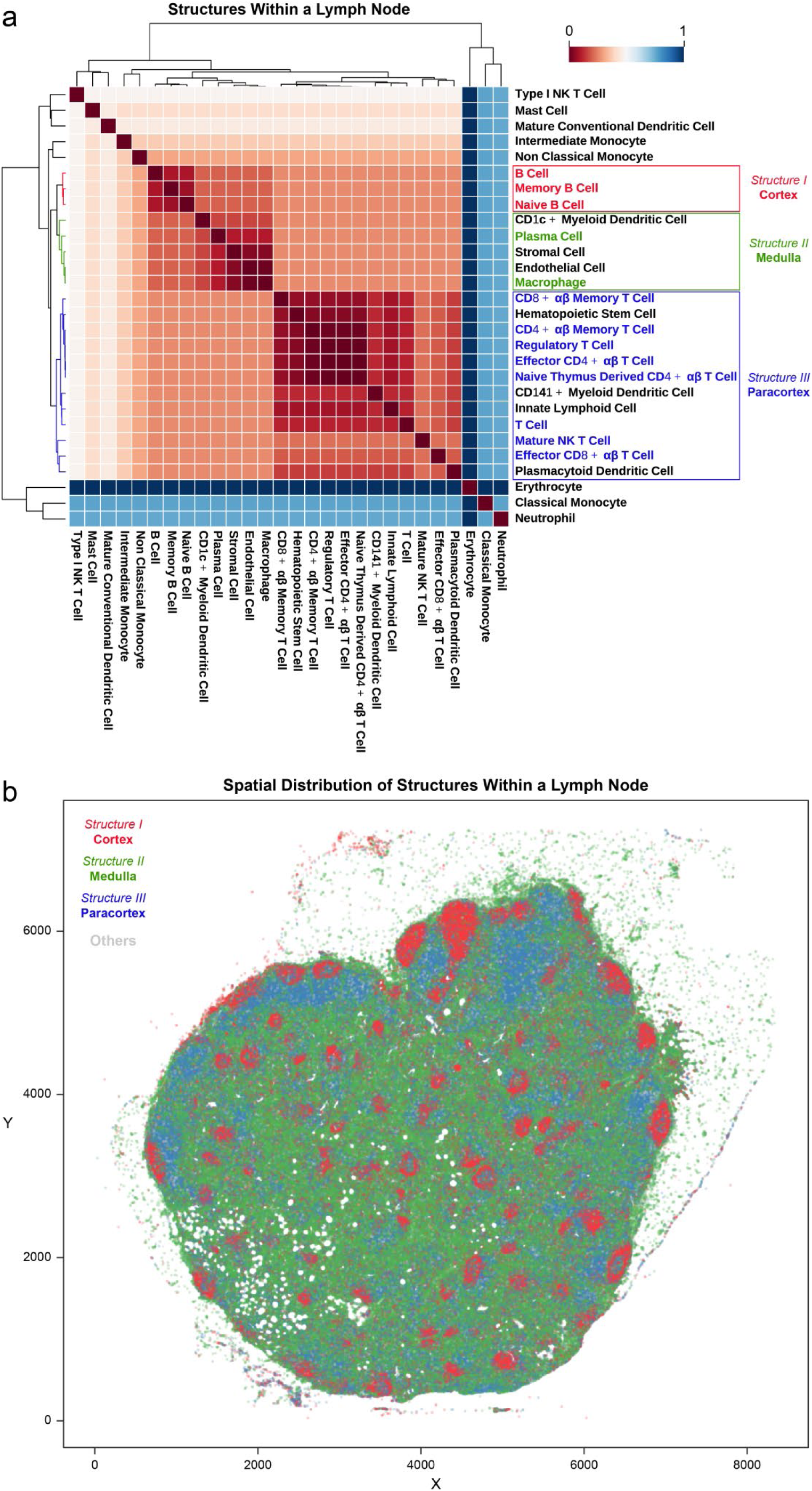
COSTE recapitulates known spatial substructures in a human lymph node. (a) SSS heatmap of immune and stromal cell types in a human lymph node highlights three major spatial modules corresponding to the lymph node’s anatomical compartments. Structure I (red) consists of B cell-enriched cortical areas (e.g., naive and memory B cells, plasma cells), Structure II (green) represents the medullary region (mix of plasma cells, macrophages, stromal and endothelial cells), and Structure III (blue) corresponds to the T cell-rich paracortex (e.g., CD4+ and CD8+ T cells). (b) The spatial distribution of these identified structures aligns with lymph node histology: B cell follicles (Structure I) appear as discrete circular clusters embedded within the medulla and are spatially segregated from the paracortical T cell zones (Structure III). The medullary region (Structure II) surrounds and supports these organized cortical and paracortical niches. This example shows that COSTE can recover known lymph-node spatial organization from spatial proximity profiles.

**Supplementary Figure 11.**
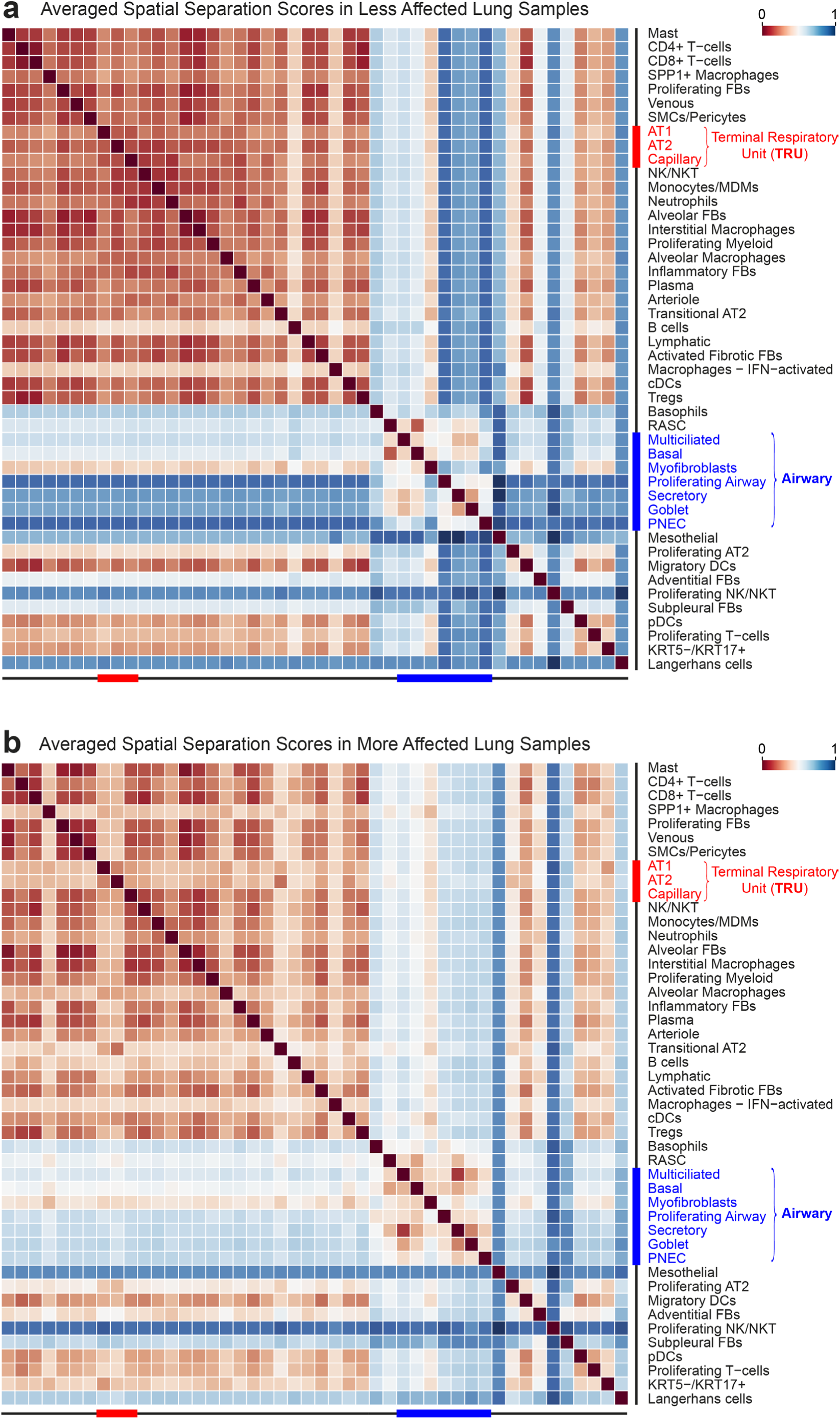
Unclustered SSS heatmaps reveal loss of normal lung structure with fibrosis progression. Unclustered SSS heatmaps from a mildly fibrotic lung sample (**a**) and a severely fibrotic lung sample (**b**) are shown with cell types ordered according to the healthy lung hierarchy. In the less affected lung, some structured pattern is still evident, whereas in the more affected lung the heatmap appears markedly disorganized. Notably, the severely fibrotic sample shows elevated SSS (weaker co-localization) among airway-related cell types, indicating that the normally cohesive airway–TRU arrangement has broken down. Overall, higher spatial separation and a loss of clear block patterns in the more affected lung reflect the dispersion of both airway and alveolar cell populations as fibrosis advances, in contrast to the orderly compartmentalization seen in healthy lungs.

**Supplementary Figure 12.**
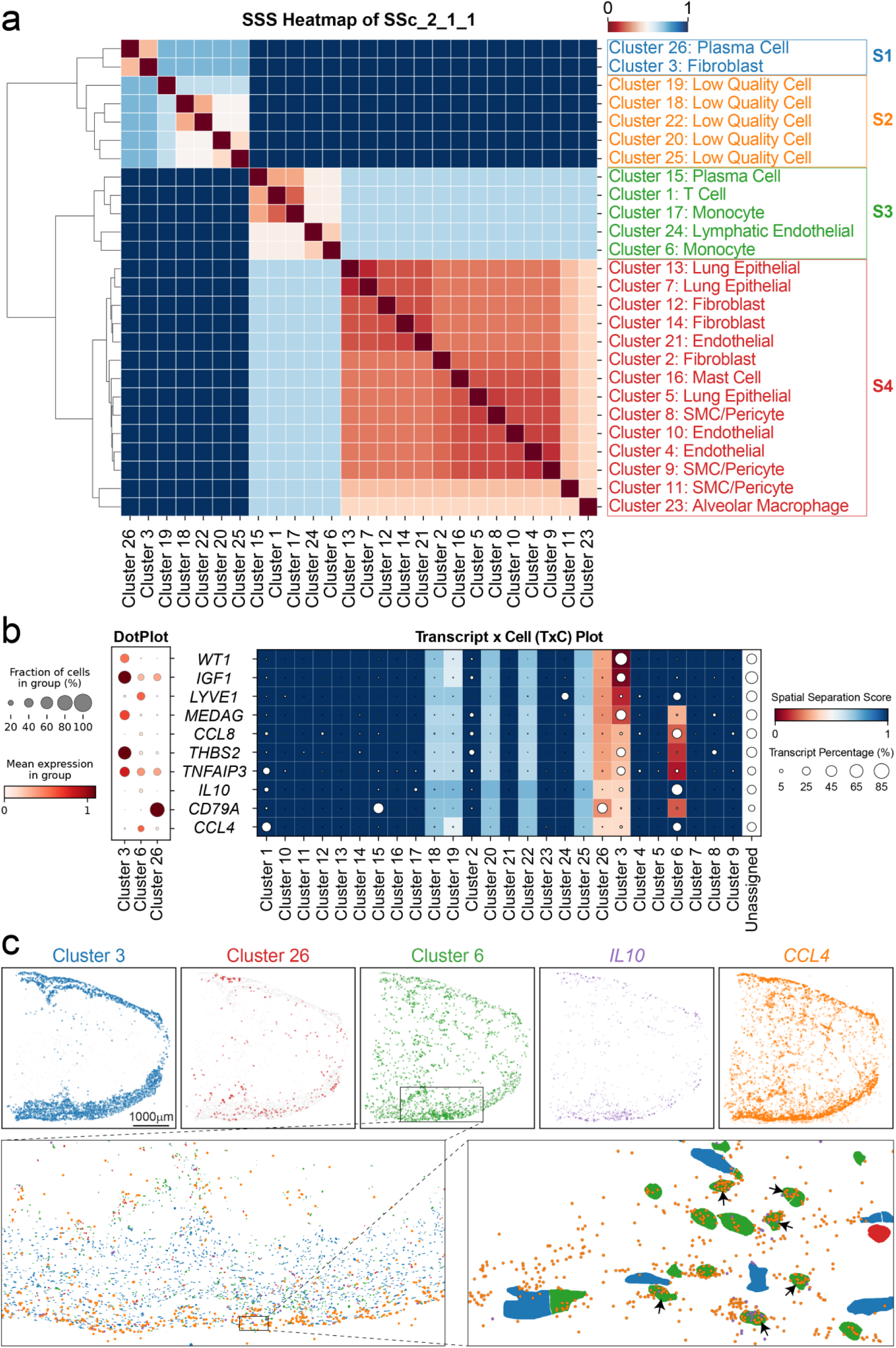
Transcript-level COSTE analysis nominates a pleura-associated monocyte state in fibrosis. (a) An SSS heatmap among 26 annotated cell clusters in one systemic sclerosis lung sample reveals four spatial structures. Structure 1 consists of fibroblasts (cluster 3) and plasma cells (cluster 26), corresponding to the fibrotic pleural thickening region (see Fig. 3a). (b) To identify transcripts enriched in this fibroblast-enriched niche, each gene was evaluated for its spatial proximity to the fibroblast cluster. A dot plot of the top ten transcripts most spatially connected to fibroblasts (cluster 3) shows that two cytokines - IL10 and CCL4 - rank highly yet are not produced by fibroblasts. Instead, IL10 and CCL4 are expressed by monocytes/macrophages (cluster 6), as indicated by the dot sizes (representing the proportion of each gene’s expression coming from each cluster). The accompanying heatmap (right) visualizes the spatial association strength of each top gene to each cluster, indicating that IL10 and CCL4 are spatially linked to the fibroblast-rich pleural structure in this sample. (c) Spatial mapping of IL10+CCL4+ cells (arrows) localizes these monocytes near the pleural boundary adjacent to fibroblasts, nominating a candidate IL10+CCL4+ pleura-associated monocyte state for follow-up validation.

**Supplementary Figure 13.**
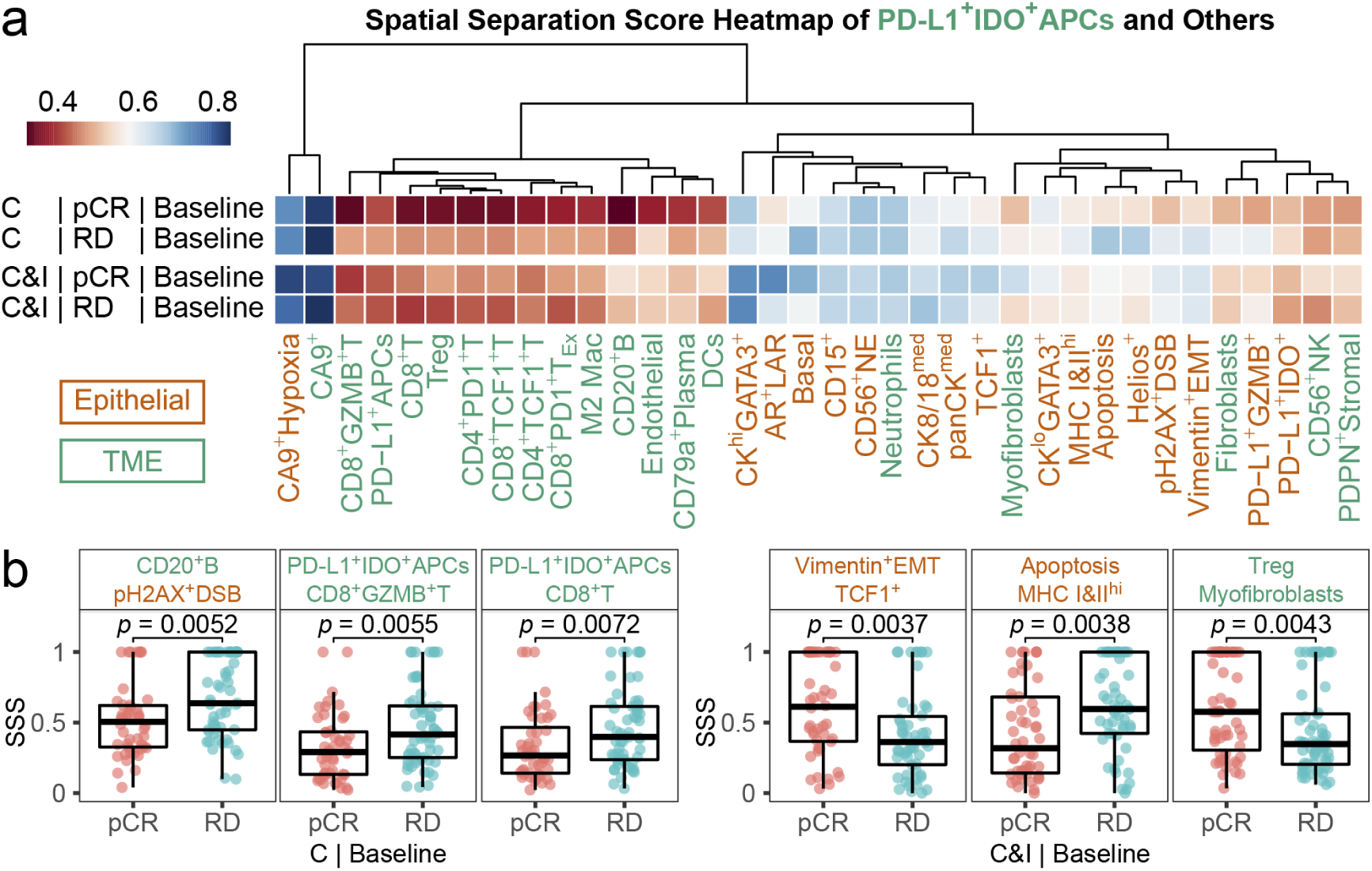
COSTE identifies response-associated immune spatial patterns in a TNBC cohort. COSTE was applied to baseline tumor biopsies from a triple-negative breast cancer (TNBC) trial to compare spatial cell-cell organization between patients with pathologic complete response (pCR) and those with residual disease (RD) under two treatments: chemotherapy (C) versus chemo-immunotherapy (C&I). (a) A heatmap of SSS values indicates that responders exhibit closer relative spatial proximity between antigen-presenting cells (PD-L1+IDO+ APCs) and other tumor or immune cell populations, whereas non-responders show greater separation between these compartments. (b) Quantification of selected cell-pair separations highlights response-associated differences. For example, in the C arm, RD tumors have higher SSS (weaker co-localization) between CD20+ B cells and pH2AX+ DNA-damaged tumor cells, as well as between PD-L1+IDO+ APCs and CD8+ T cells, compared to pCR tumors. In the C&I arm, RD is associated with increased separation between mesenchymal cells undergoing EMT (Vimentin+) and TCF1+ T cells, and between regulatory T cells and myofibroblasts, relative to responders. These findings suggest that immune-stromal spatial disconnection is associated with treatment outcome in this retrospective cohort.

**Supplementary Figure 14.**
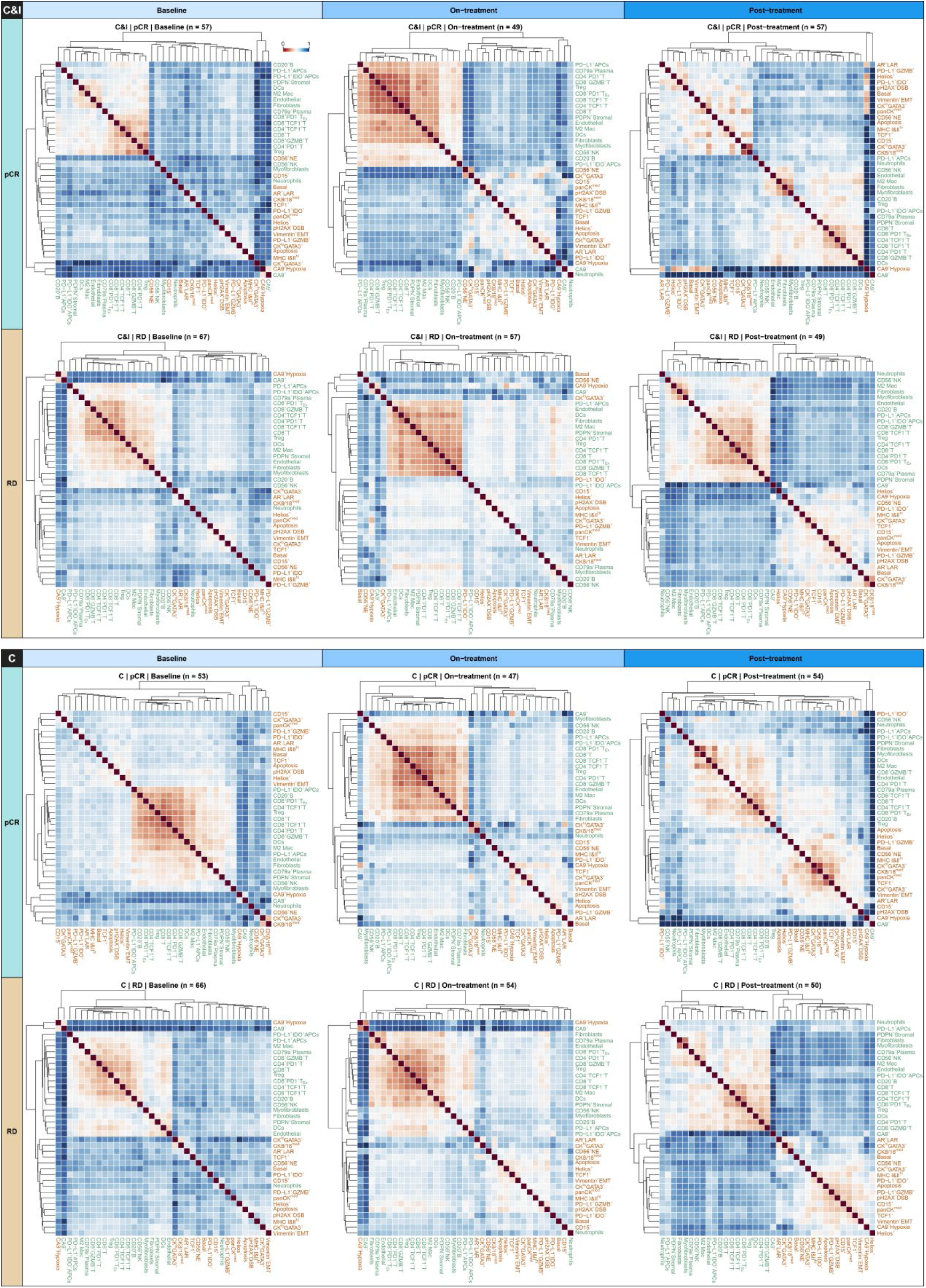
Therapy-associated spatial reorganization of tumor and microenvironment in TNBC. COSTE SSS heatmaps are presented for all 12 patient subgroups defined by treatment arm (C vs C&I), response (pCR vs RD), and time point (baseline, on-treatment, post-treatment). These heatmaps provide an overview of the dynamic spatial reconfiguration of epithelial (orange labels), immune, and stromal (green labels) compartments over the course of therapy. Changes in clustering patterns and SSS values across time indicate how different cell-cell associations strengthen or weaken with treatment. This panel highlights that treatment modality and response status are associated with distinct trajectories of spatial remodeling in the tumor microenvironment.

## Supplementary Tables

**Supplementary Table 1. Runtime and memory performance of spatial neighborhood analysis methods on the Xenium neonatal mouse pup dataset**

**Supplementary Table 2. Transcript by cell spatial separation scores in lung fibrosis from a systemic sclerosis patient**

**Supplementary Table 3. Transcript spatial separation scores in lung fibrosis from a systemic sclerosis patient using landmark clusters**

