## Supplementary figures and images for "Cophenetic Spatial Topology Embedding reveals multiscale tissue architecture in spatial omics"

### Supplementary Fig. 1

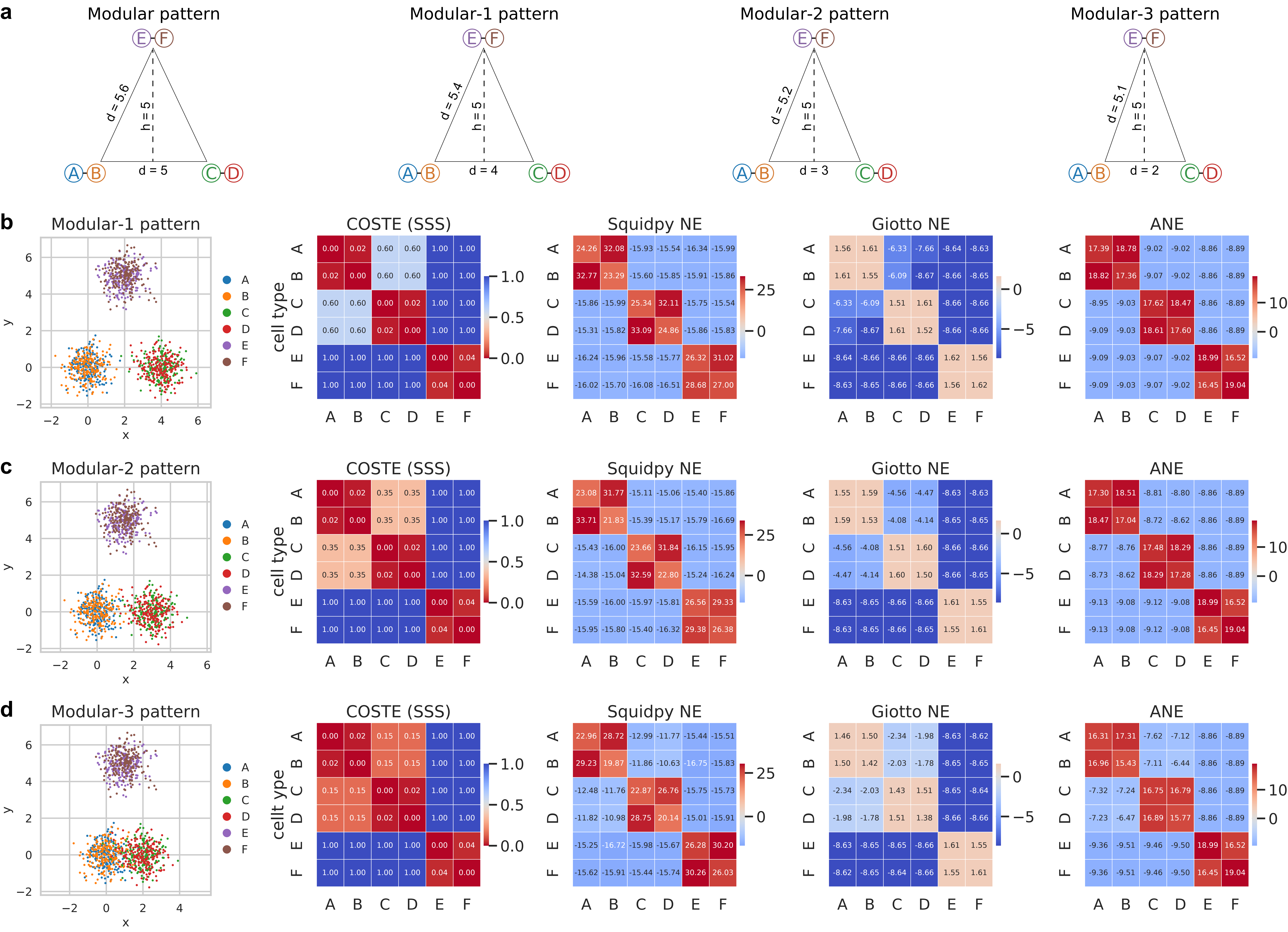

### Supplementary Fig. 2

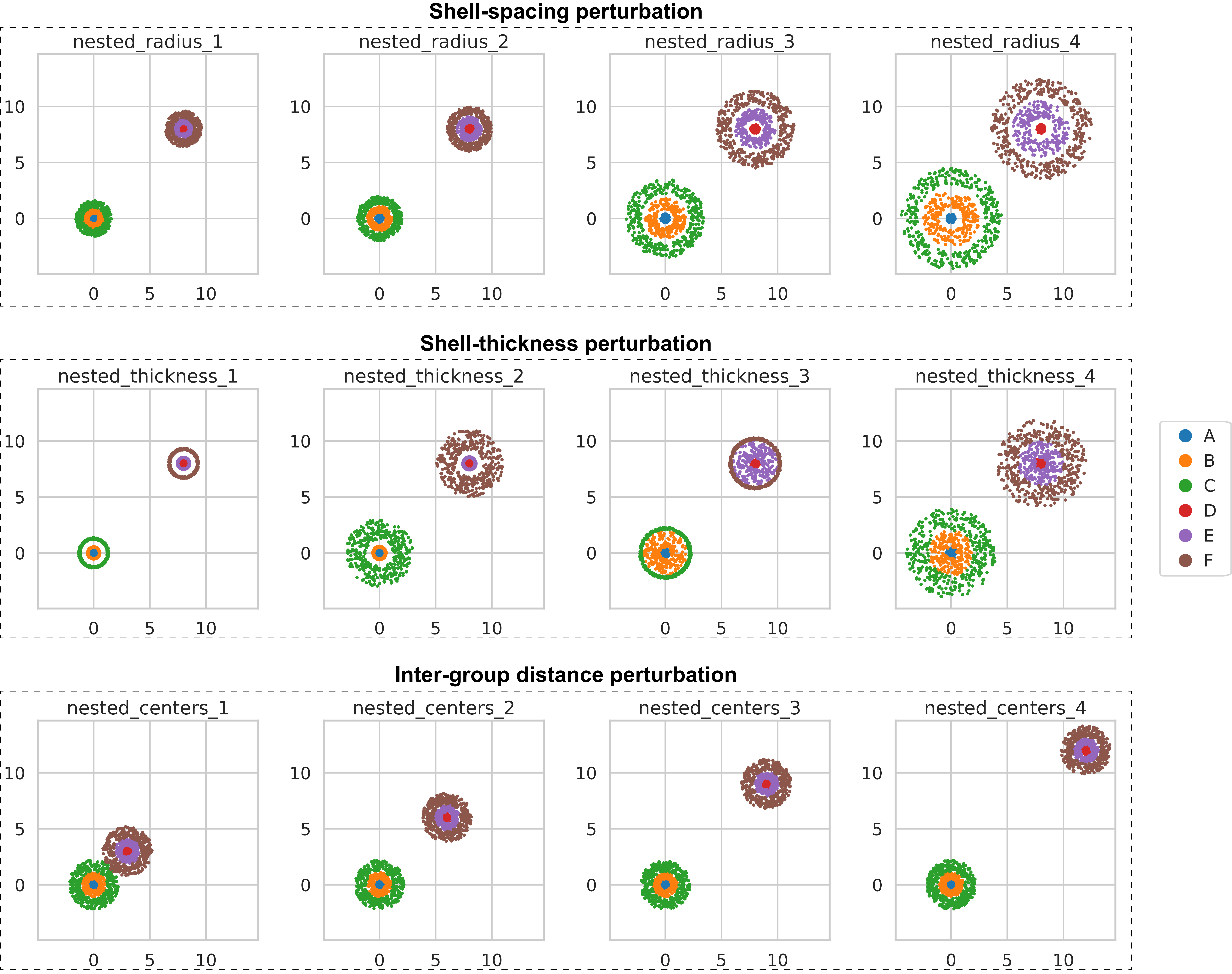

### Supplementary Fig. 3

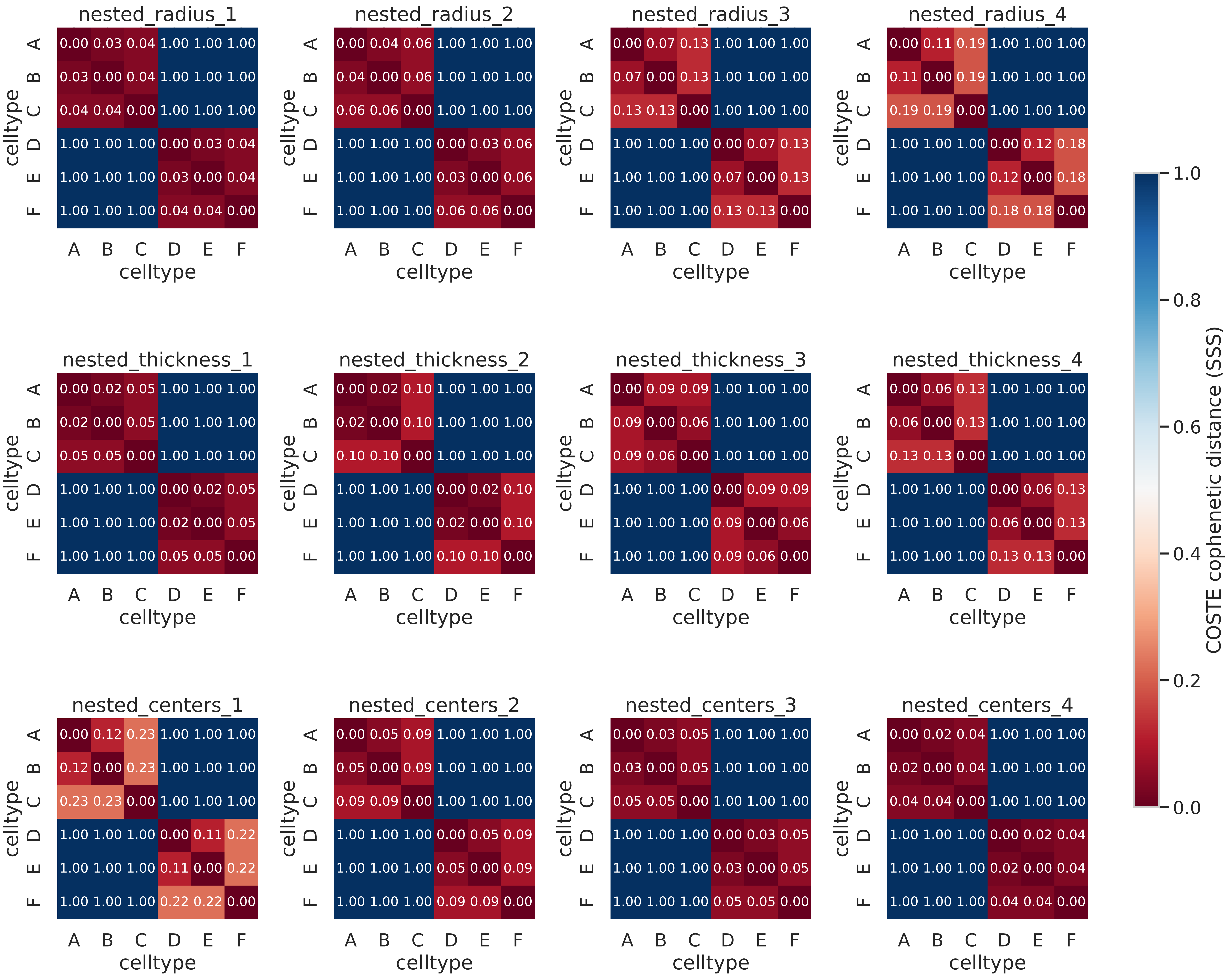

### Supplementary Fig. 4

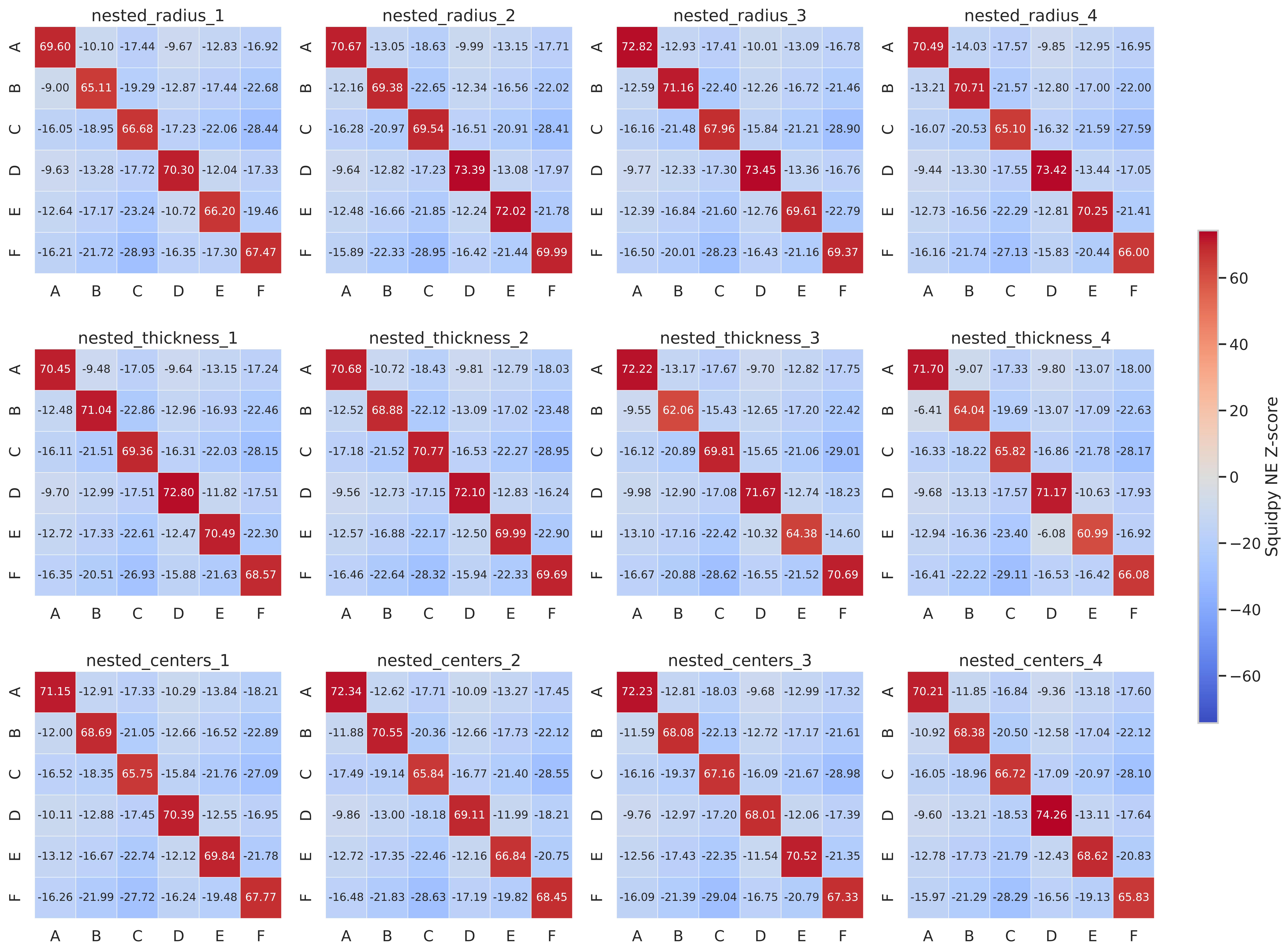

### Supplementary Fig. 5

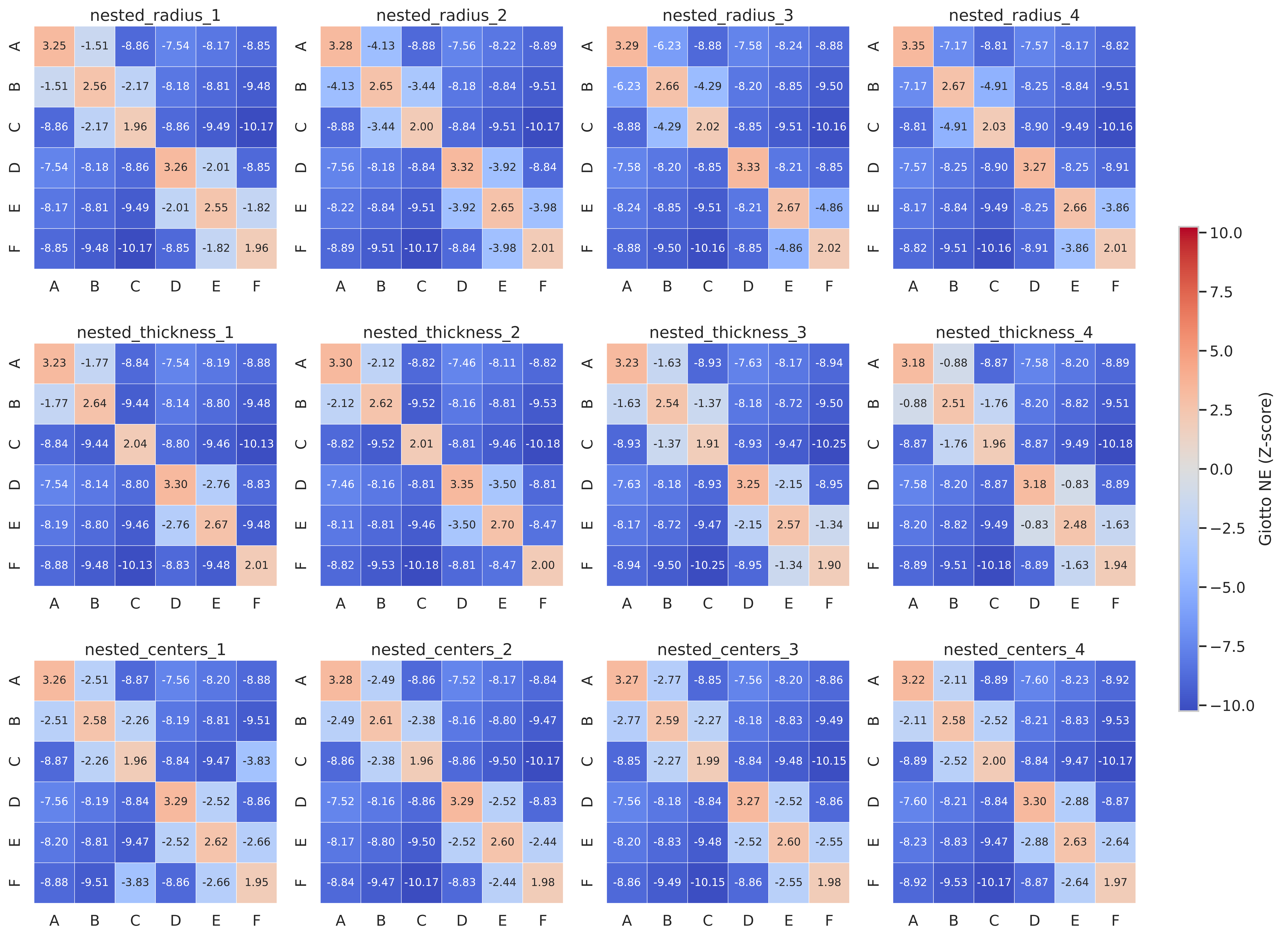

### Supplementary Fig. 6

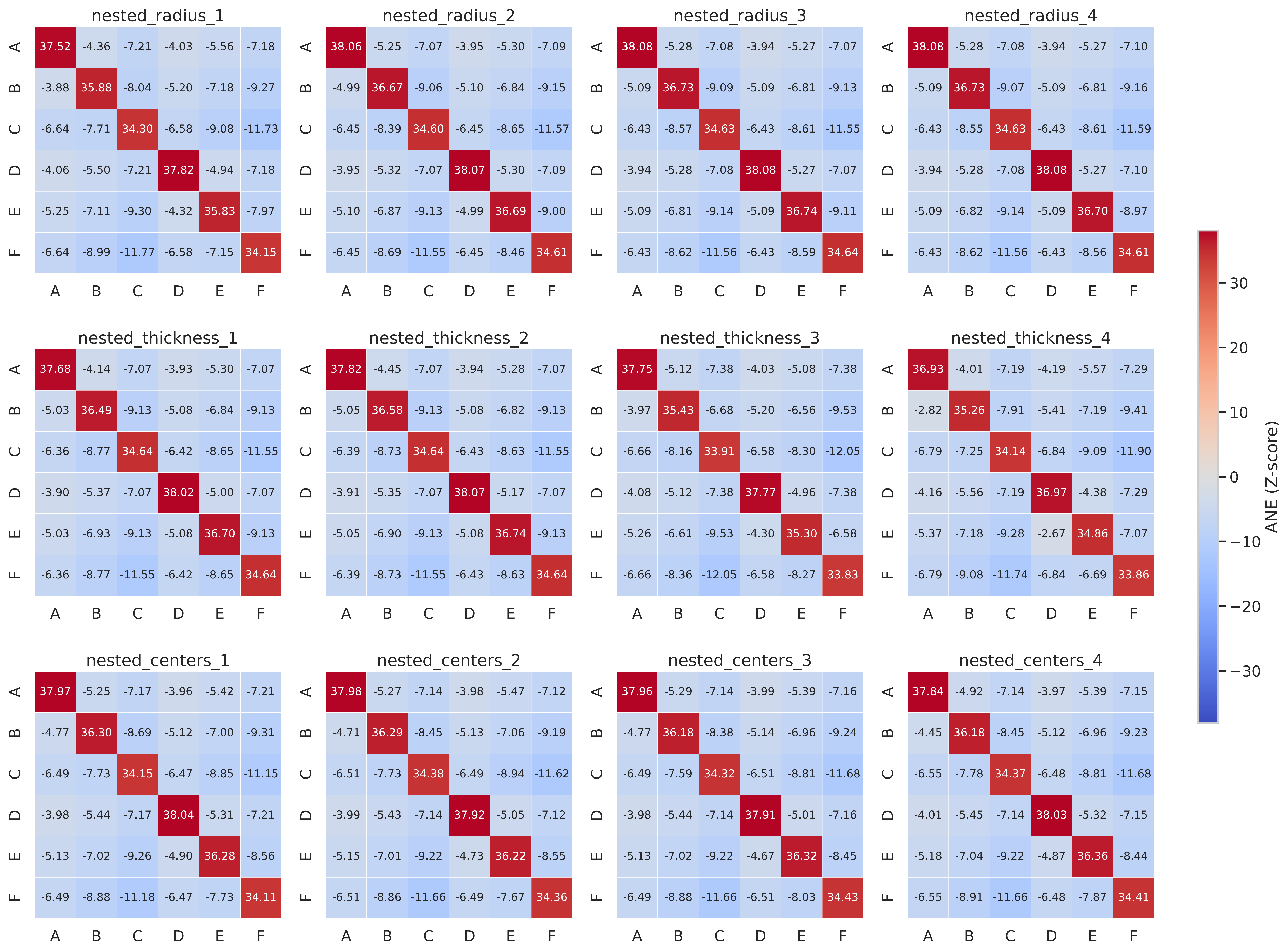

### Supplementary Fig. 7

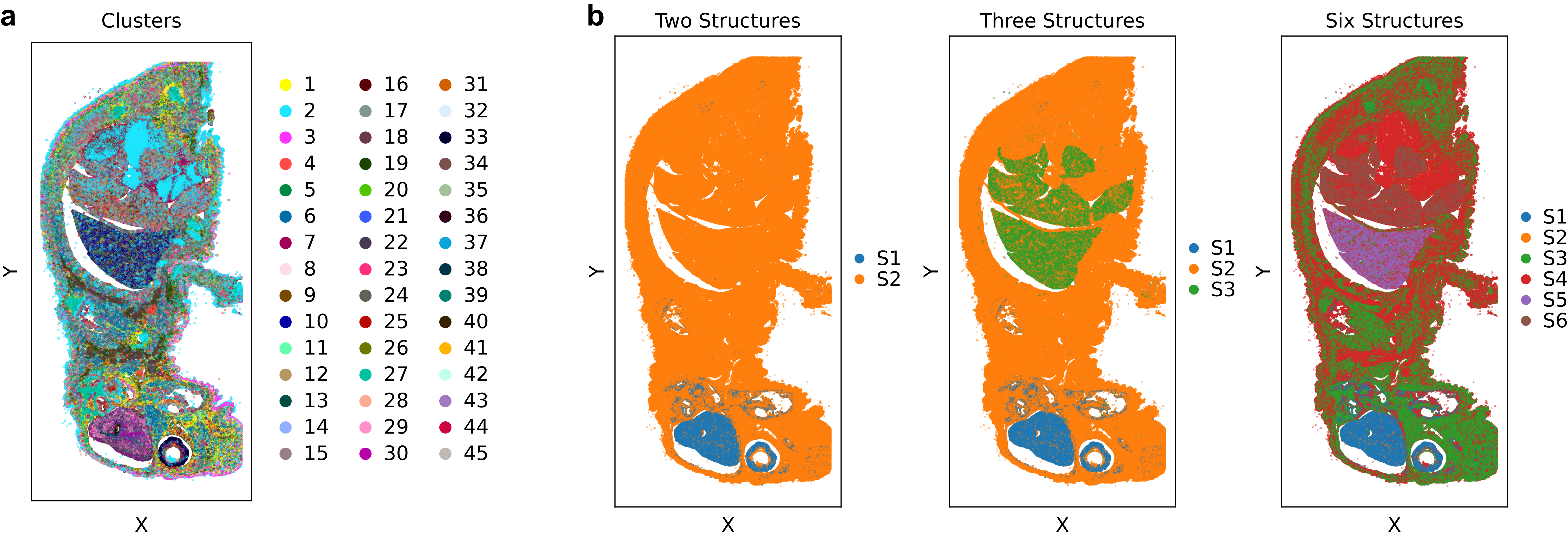

### Supplementary Fig. 8

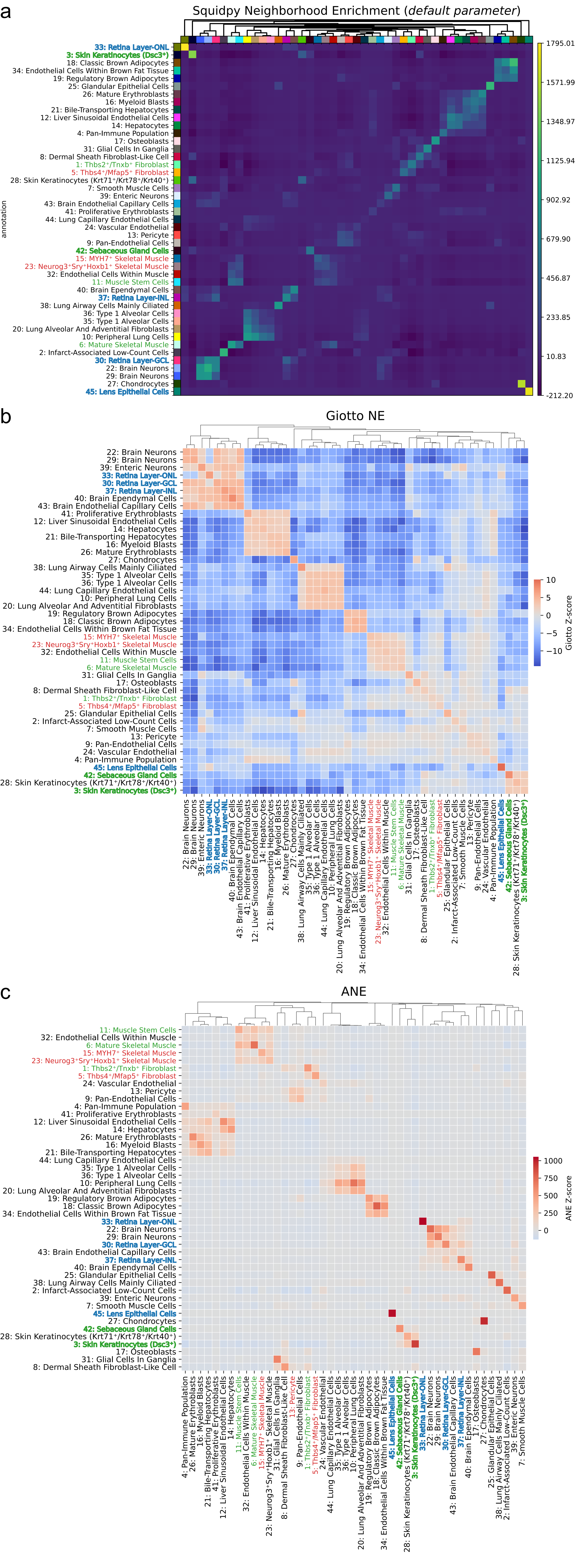

### Supplementary Fig. 9

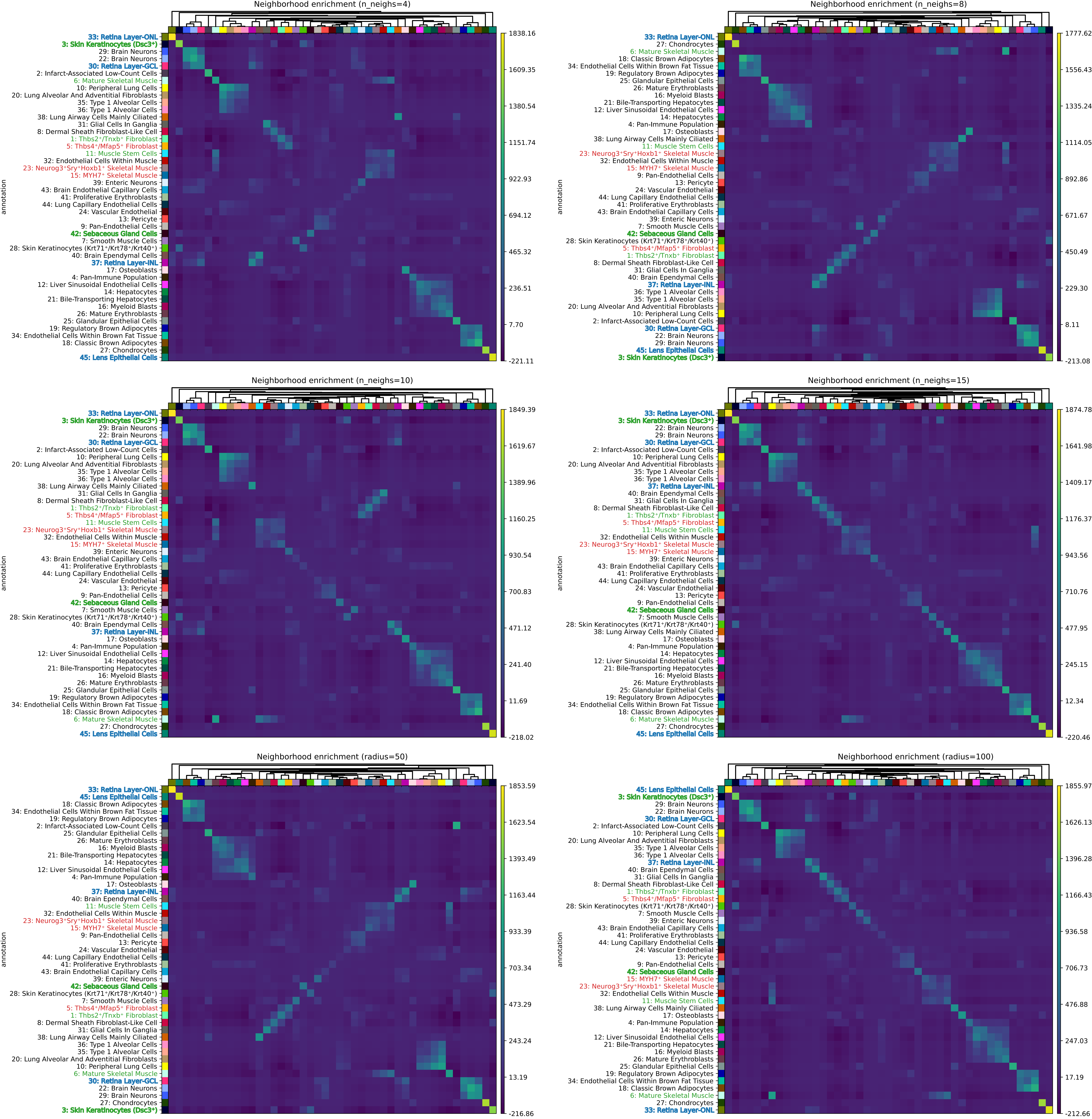

### Supplementary Fig. 10

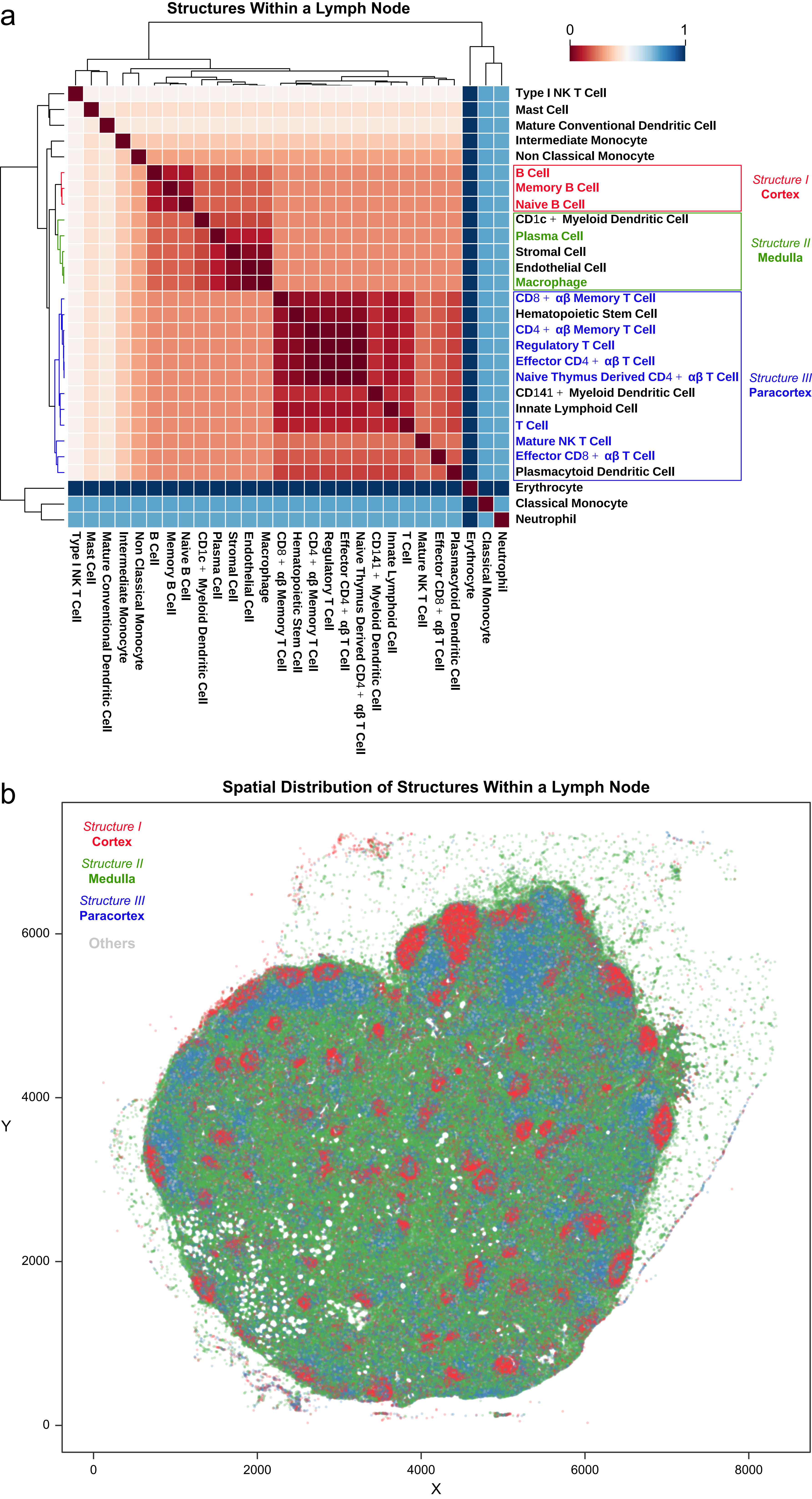

### Supplementary Fig. 12

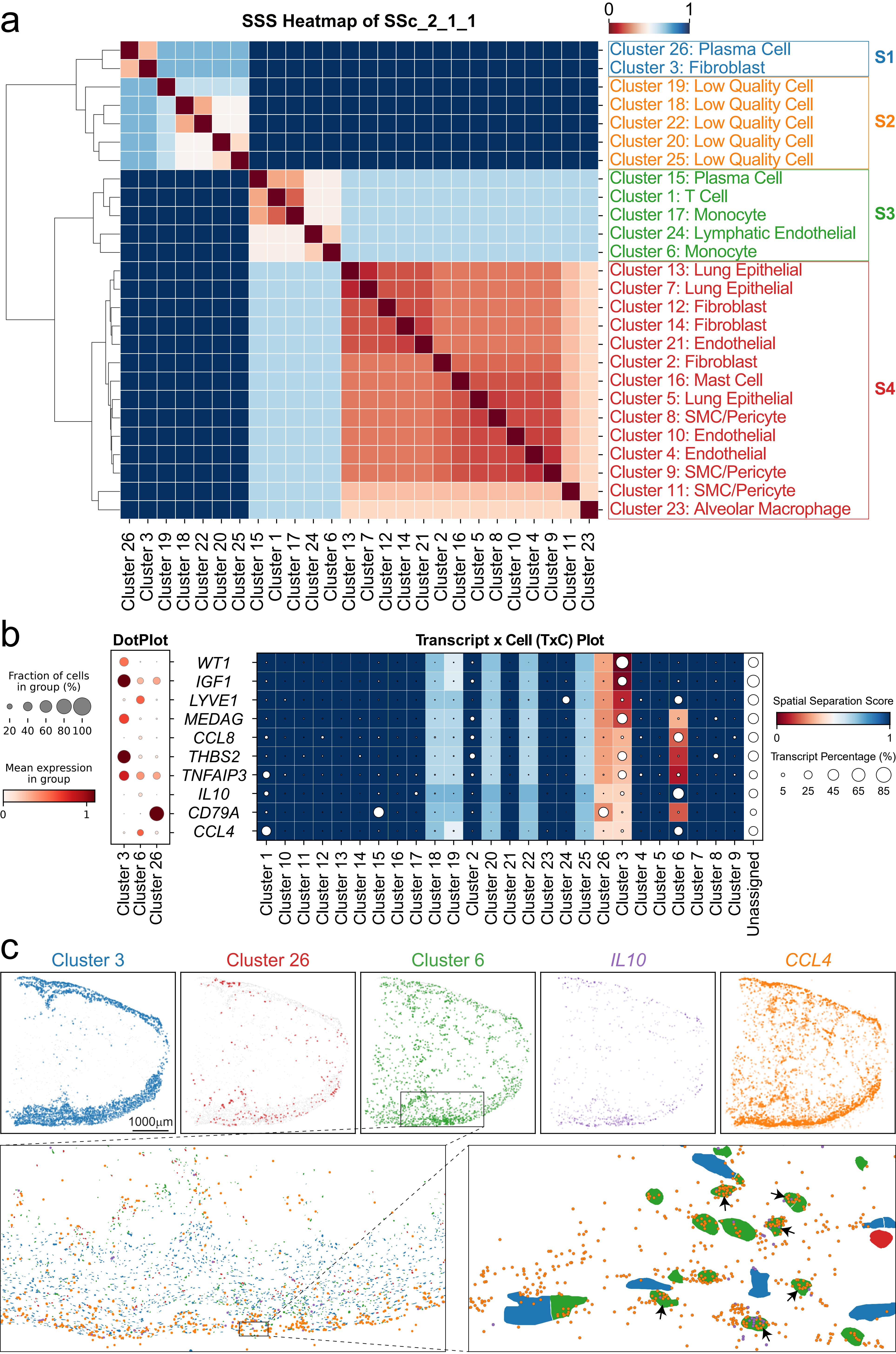

### Supplementary Fig. 13

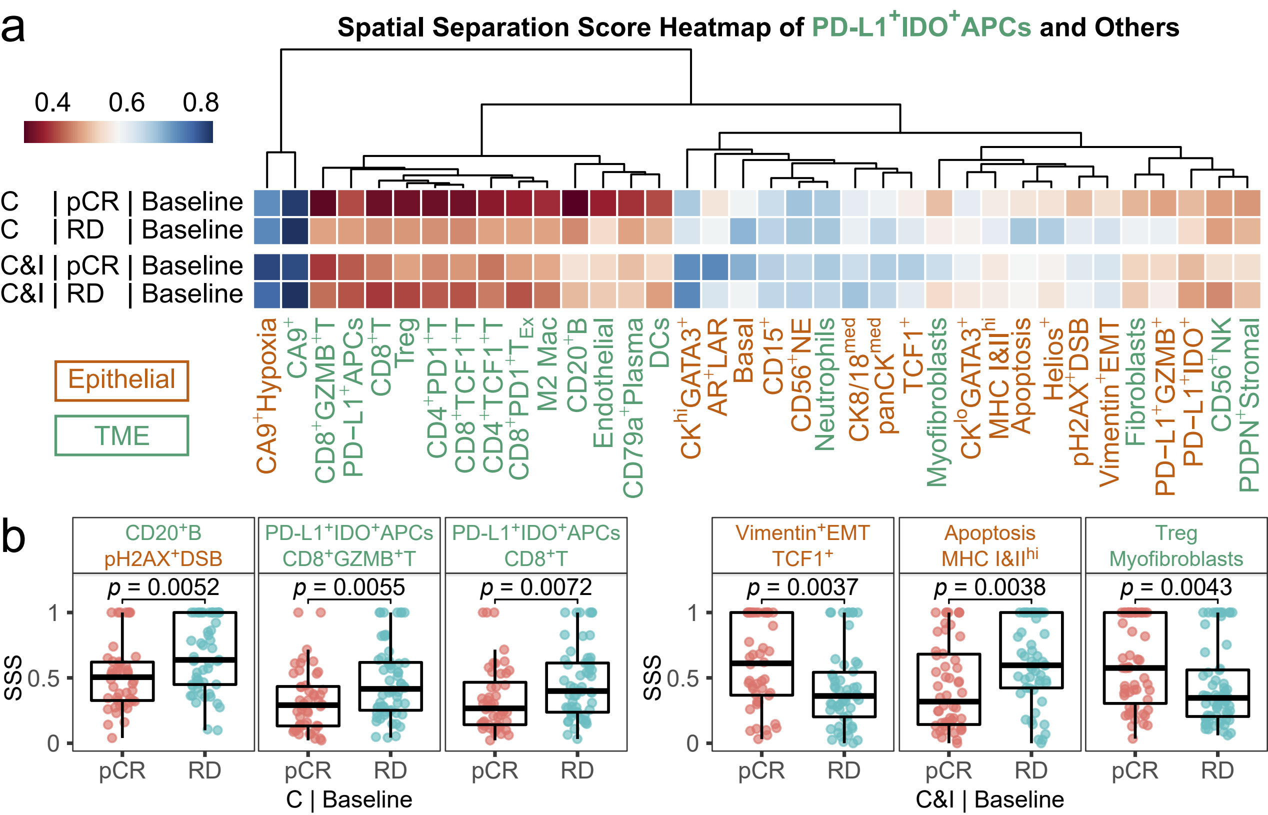

### Supplementary Fig. 14

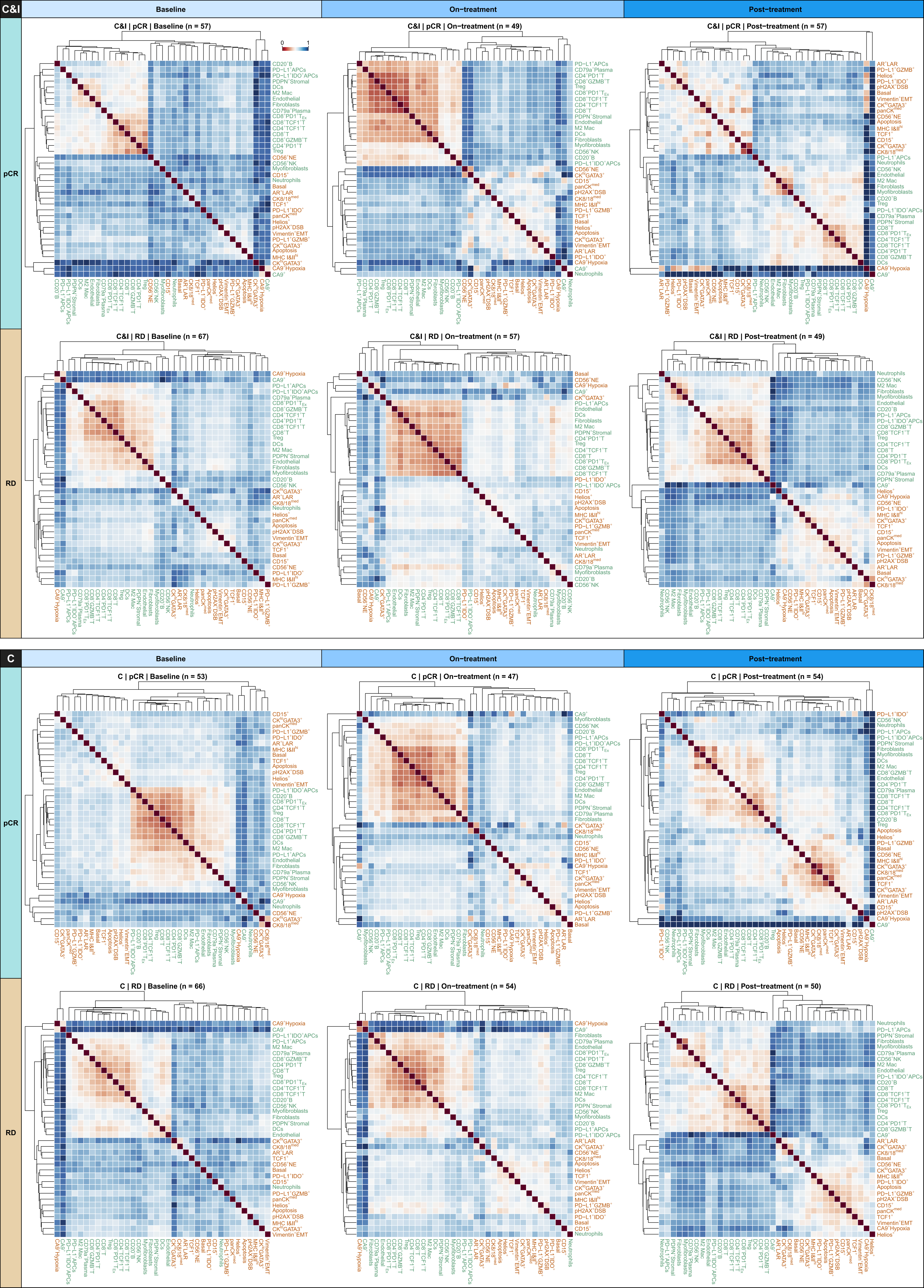
