## Supplementary Fig. 11 for "Cophenetic Spatial Topology Embedding reveals multiscale tissue architecture in spatial omics"

**a** Averaged Spatial Separation Scores in Less Affected Lung Samples

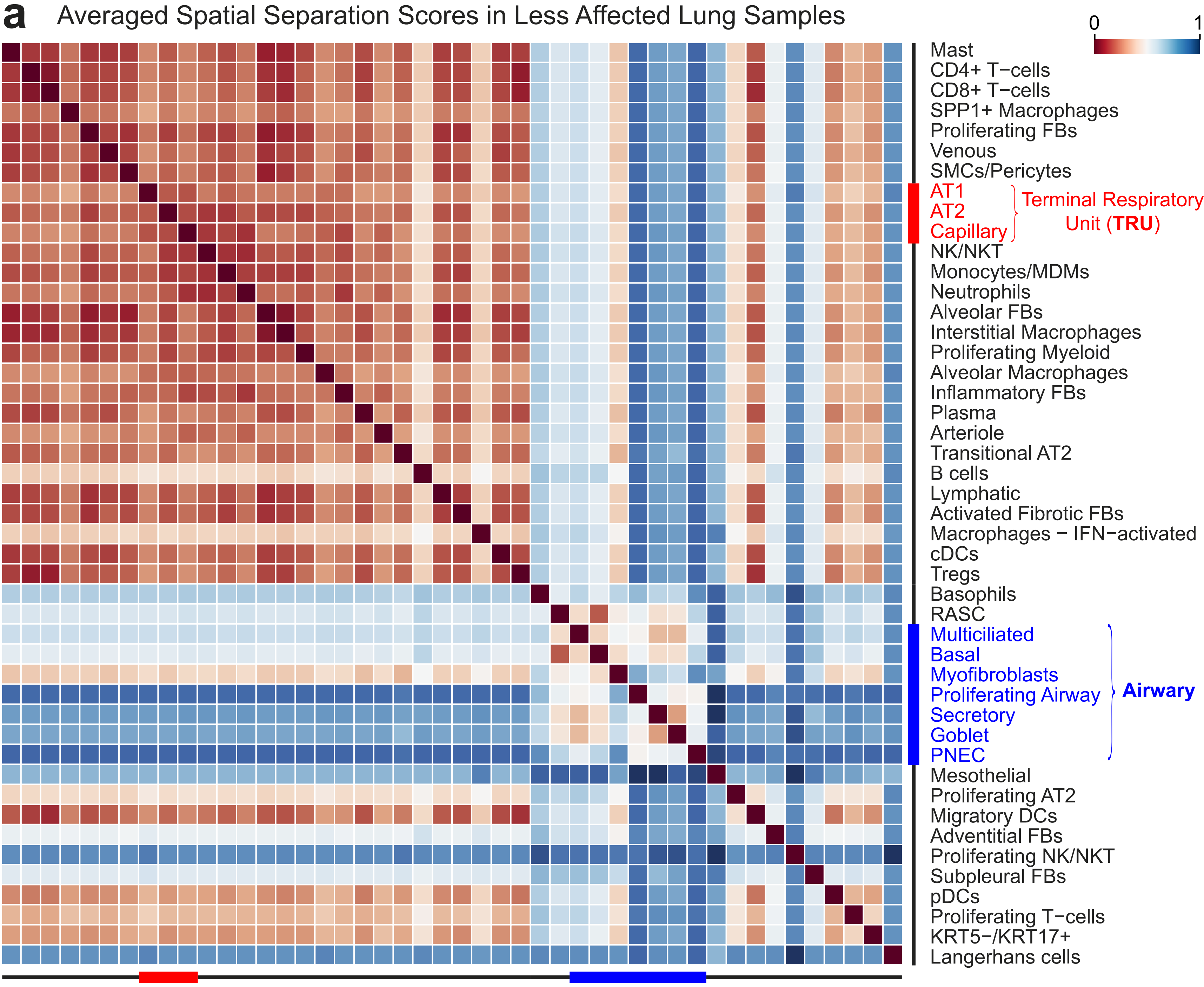


**b** Averaged Spatial Separation Scores in More Affected Lung Samples

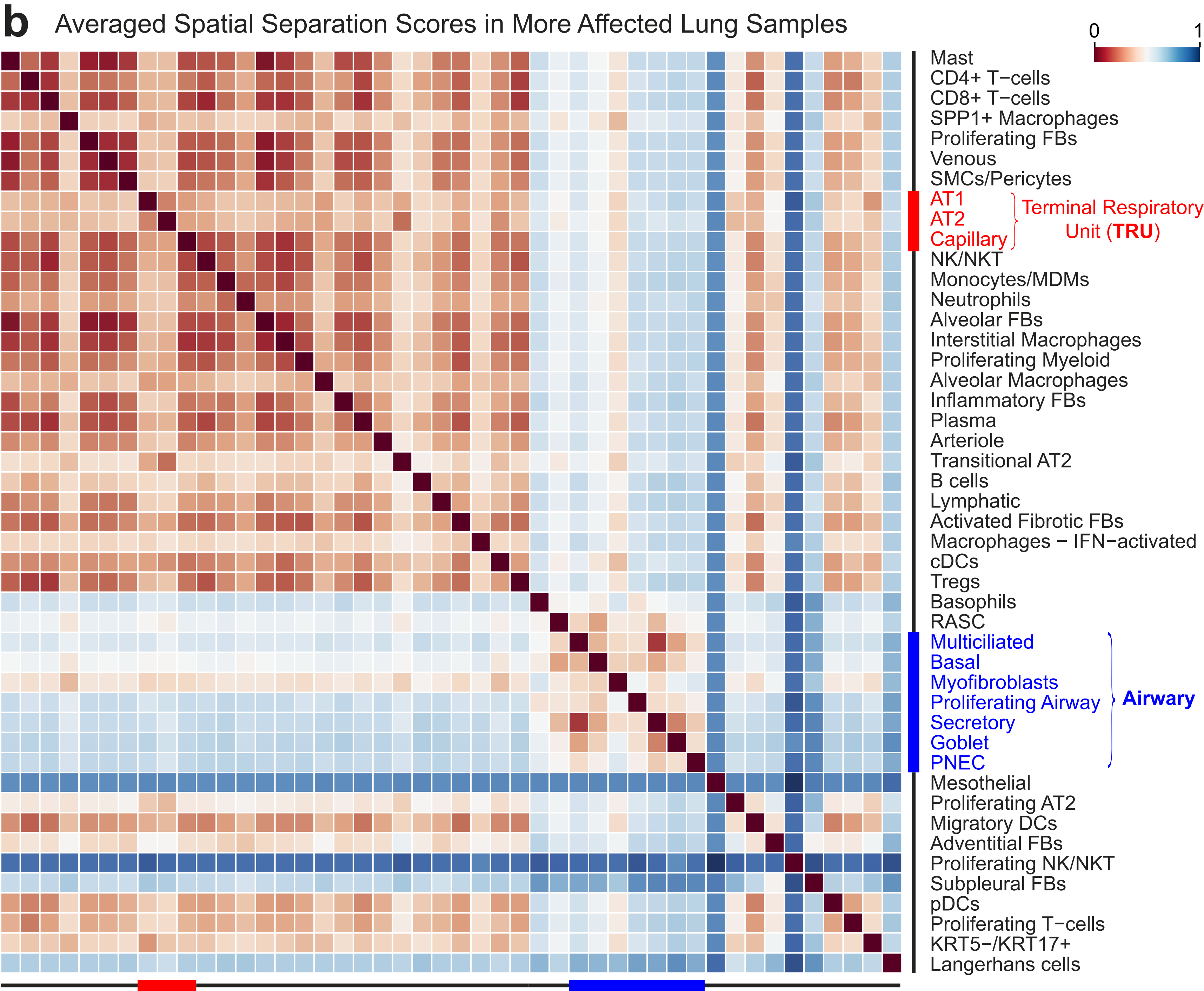
