## Supplementary Table 1 for "Cophenetic Spatial Topology Embedding reveals multiscale tissue architecture in spatial omics"

**Supplementary Table 1** | Runtime and memory performance of spatial neighborhood analysis methods on the Xenium neonatal mouse pup dataset.

| Method | Number of Permutations (if applicable) | Runtime (seconds) | Peak Memory (GB) | Minor Page Faults | Context Switches | Throughput (cells/sec) |
| --- | --- | --- | --- | --- | --- | --- |
| COSTE | – | 74.9 | 8.92 | 100,332 | 1,114 | 17,311 |
| Squidpy | 1000 | 248.3 | 5.20 | 57,932,245 | 92,604 | 5,221 |
| Giotto | 100 ^*^ | 1031 | 38.64 | 128,890,561 | 54,568 | 1,257 |
| ANE | – | 2.1 | 43.59 | 202,256 | 93 | 620,234 |

**Runtime:** wall time from method start to completion in seconds.

**Peak Memory:** maximum resident set size (RSS) during execution (in gigabytes).

**Minor Page Faults:** number of memory address translation faults handled without requiring disk I/O.

**Context Switches:** total count of voluntary plus involuntary context switches incurred, as reported by the operating system.

**Throughput:** total number of cells processed divided by runtime (cells per second).

^*^ For Giotto, the number of permutations was set to 100 instead of the default 1,000 because the default setting caused out‑of‑memory errors on the Xenium neonatal mouse pup dataset. This reduction does not affect the qualitative comparison of enrichment patterns but substantially improves runtime and memory stability on this large dataset.
